# Mitochondrial network reorganization and transient expansion during oligodendrocyte generation

**DOI:** 10.1101/2023.12.05.570104

**Authors:** Xhoela Bame, Robert A. Hill

## Abstract

Oligodendrocyte precursor cells (OPCs) give rise to myelinating oligodendrocytes of the central nervous system. This process persists throughout life and is essential for recovery from neurodegeneration. To better understand the cellular checkpoints that occur during oligodendrogenesis, we determined the mitochondrial distribution and morphometrics across the oligodendrocyte lineage in mouse and human cerebral cortex. During oligodendrocyte generation, mitochondrial content expanded concurrently with a change in subcellular partitioning towards the distal processes. These changes were followed by an abrupt loss of mitochondria in the oligodendrocyte processes and myelin, coinciding with sheath compaction. This reorganization and extensive expansion and depletion took 3 days. Oligodendrocyte mitochondria were stationary over days while OPC mitochondrial motility was modulated by animal arousal state within minutes. Aged OPCs also displayed decreased mitochondrial size, content, and motility. Thus, mitochondrial dynamics are linked to oligodendrocyte generation, dynamically modified by their local microenvironment, and altered in the aging brain.

## INTRODUCTION

Insulating, lipid-rich myelin sheaths enwrap axons to accelerate electrical transmission and provide trophic support to neurons^1,2^. The myelin of the central nervous system is synthesized and maintained by oligodendrocytes which derive from a resident population of oligodendrocyte precursor cells (OPCs). Oligodendrocyte generation persists throughout life holding important roles for plasticity, cognition, and myelin replacement in neurodegenerative pathologies^2,3^. The developmental stages of oligodendrocytes consist of migration and proliferation of OPCs, differentiation, and maturation into myelinating oligodendrocytes^4^. Several intrinsic and extrinsic cues, involving transcription and growth factors, axon diameter, and neuronal activity can modulate each of these stages^5–8^. However, the precise mechanisms that guide OPCs to differentiate or undergo cell death are not well defined. To better understand this process, the physiological and subcellular events that occur within these cells in their native environment must be determined.

Mitochondrial activity drives the differentiation, maturation, and cell death of many cell types throughout the body^9–18^. Genes linked to these mitochondrial dynamics vary across the oligodendrocyte lineage^19–21^ suggesting that mitochondrial activity could be correlated with fate decisions in this lineage. This is supported by changes in mitochondria during the differentiation of cultured oligodendrocytes^22^ along with the association of mitochondria-regulated signaling pathways in modulating OPC survival and myelination^23,24^. Moreover, mitochondria motility in oligodendrocytes appears low^25,26^ but whether the same dynamics apply to OPCs has not been reported. Finally, disrupted mitochondrial function have been associated with aging and neurodegeneration^27,28^. Indeed, aged OPCs, characterized by their reduced potential to differentiate, exhibit hallmarks of mitochondrial dysfunction and restoring mitochondrial function in these cells can partially restore their ability to differentiate^29^.

These observations position mitochondria as multifunctional, plastic organelles with the potential to regulate the fate of OPCs. However, there has never been a systematic analysis of mitochondrial dynamics as OPCs are self-renewing, differentiating, or dying. Here, we used a combination of intravital fluorescence and label-free optical imaging to determine the mitochondrial dynamics during these fate decisions in the live intact mouse brain. This was coupled with ultrastructural 3-dimensional analysis of mitochondria morphometrics in human tissue. Mitochondria content changed drastically during oligodendrocyte generation with myelinating oligodendrocytes exhibiting the lowest mitochondrial content out of all stages. Mitochondria morphology transitioned from more elongated in OPCs to fragmented in myelinating oligodendrocytes. Longitudinal imaging as an OPC transitioned into a myelinating oligodendrocyte revealed a migratory pattern of mitochondria out of the soma and into the processes and the newly formed myelin sheaths. This was followed by an abrupt mitochondrial loss as the oligodendrocyte matured and the myelin sheaths compacted. Once the oligodendrocytes matured, their mitochondria were remarkably stable over days. In contrast, OPC mitochondria were dynamic and continuously moved throughout the cell. This motility was affected by the animal arousal state as anesthesia and sedation decreased mitochondria motility. Lastly, alterations in OPC mitochondria volume fraction, size, and motility were found in aged animals. These results reveal the spatiotemporal dynamics of mitochondria at different fates and ages of OPCs and uncover new pathways that impact these dynamics in the live brain.

## RESULTS

### Labeling and intravital imaging of mitochondria in the oligodendrocyte lineage

Mitochondria are multifunctional organelles known to drive cellular fate but their involvement with oligodendrocyte generation has never been characterized in the live intact brain. To enable visualization of mitochondria specifically in the oligodendrocyte lineage, we generated a triple transgenic mouse line, *Cspg4*-creER: Ai9: PhAM, that allows for the expression of the mitochondria and cell cytoplasmic reporters (respectively, mito-Dendra2 and cyto-tdTomato) in chondroitin sulfate proteoglycan 4 (Cspg4)-expressing cells after tamoxifen-inducible Cre recombination (Fig. 1 and Fig. 2a). To have sparse labeling while also capturing cells at different stages of differentiation, a low dose of tamoxifen was injected at postnatal day (P) 25 to induce Cre recombination. Intravital imaging over the somatosensory cortex was done 3 weeks later (Fig. 1a). Thus, during imaging, there was a mixture of cells at the OPC stage, cells actively differentiating and cells that had differentiated into myelinating oligodendrocytes.

**Figure 1:**
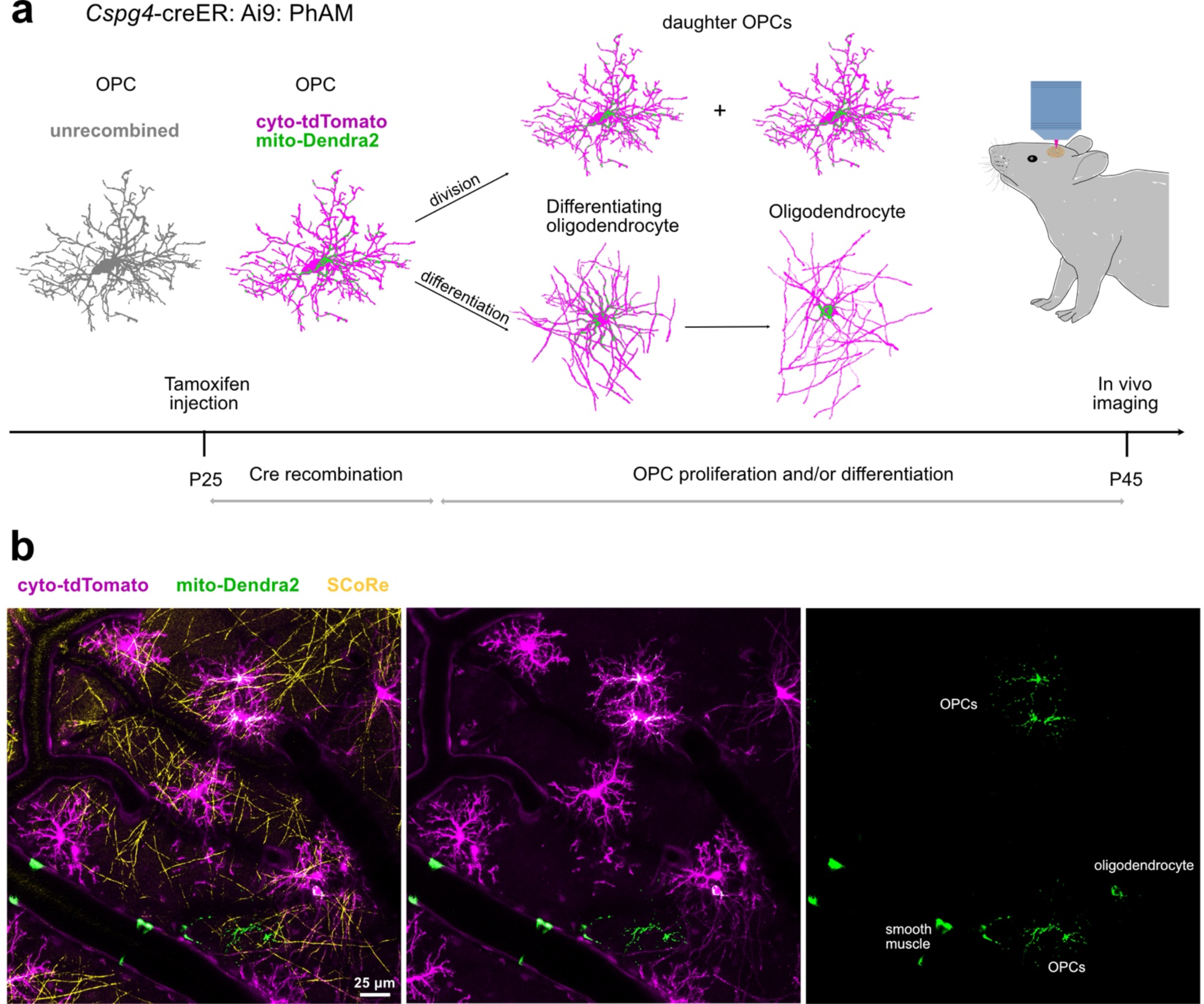
In vivo imaging of mitochondria in the oligodendrocyte lineage. **a,** Schematic illustration of experimental timeline. Tamoxifen was injected at postnatal day (P) 25 to induce Cre recombination in the *Cspg4*-creER: Ai9: PhAM triple transgenic mouse initiating labeling of cytoplasm (tdTomato) and mitochondria (Dendra2) in Cspg4-expressing cells and their progeny. Imaging was performed 20 days after in layer 1 of the somatosensory cortex. During this time, spontaneous events such as OPCs proliferation and/or differentiation into myelinating oligodendrocytes occur, populating the imaging region with recombined cells of the oligodendrocyte lineage. **b,** Representative in vivo image of the full imaging window in one position. A subset of OPCs and their progeny are either single or dual labeled with cyto-tdTomato and/or mito-Dendra2. Other Cspg4-expressing cells such as smooth muscle cells are also labeled and distinguished based on their location and morphology and excluded from the analysis. SCoRe signal comes from compact myelin sheaths in the territory.

**Figure 2:**
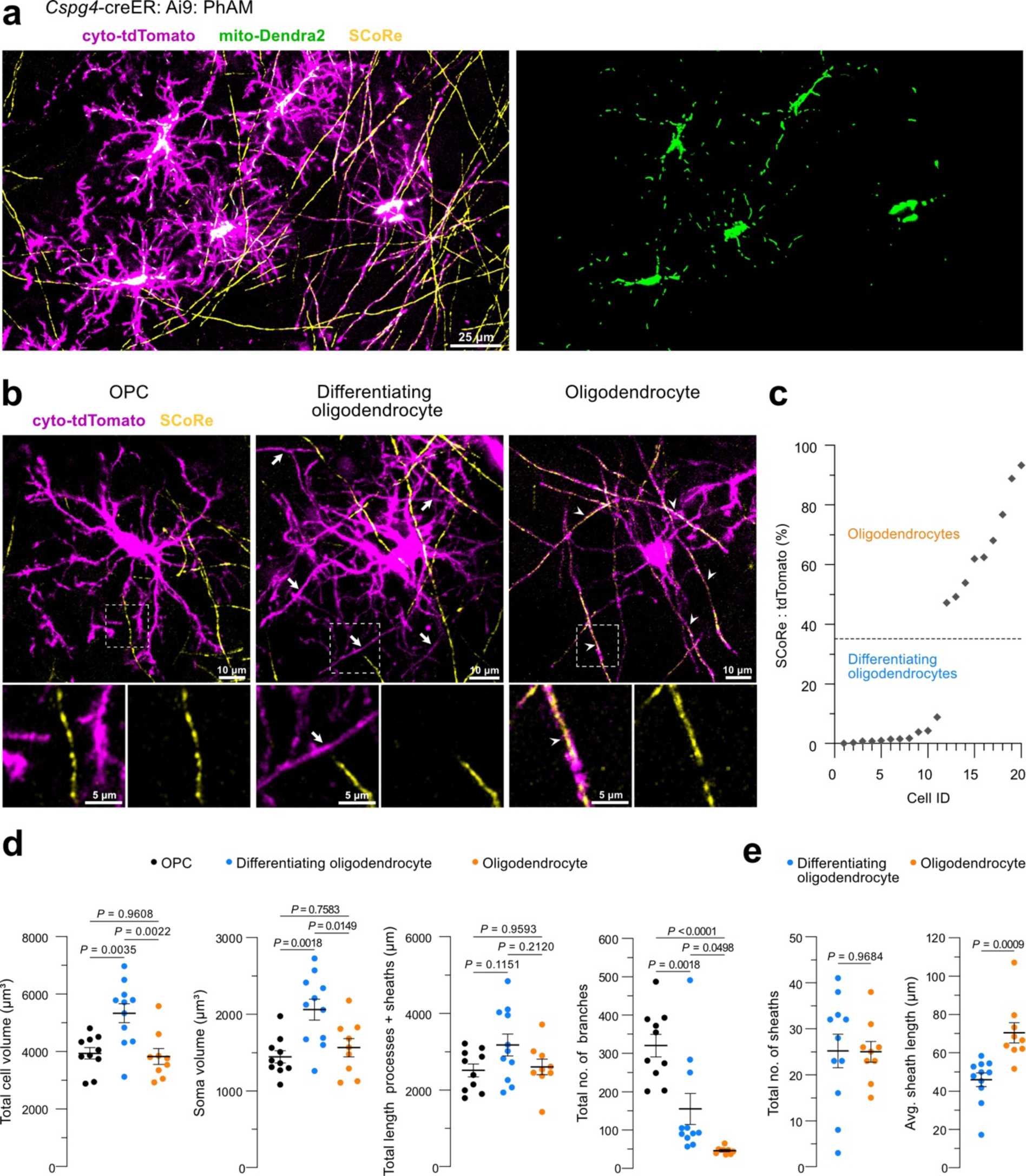
Imaging the oligodendrocyte lineage and their mitochondria in the intact cerebral cortex. **a,** In vivo image captured from the *Cspg4-*creER: Ai9: PhAM mouse cortex with labeling of mitochondria (green) in the oligodendrocyte lineage (magenta). Spectral confocal reflectance microscopy (SCoRe, yellow) reveals compact myelin sheaths. **b,** In vivo images showing the morphology of an OPC (left), differentiating oligodendrocyte (middle) and myelinating oligodendrocyte (right). Lack of SCoRe signal in the differentiating oligodendrocyte sheaths (arrows) and presence of SCoRe signal in the sheaths coming from a myelinating oligodendrocyte (arrowheads). **c,** Quantification of SCoRe coverage along the sheaths displayed as a SCoRe to cytoplasmic tdTomato ratio. Cells with high SCoRe coverage in the sheaths (>34.54%) were grouped as myelinating oligodendrocytes. Cells with low SCoRe coverage in the sheaths (<34.54%) were grouped as differentiating oligodendrocytes (*n* = 20 cells from 6 mice). **d,** Differentiating oligodendrocytes occupy the highest total cell and soma volume. Total length of processes (+ sheaths when present) is comparable among all the three stages while total number of branches per cell is highest for the OPCs (*n* = 10 OPCs, 11 differentiating and 9 myelinating oligodendrocytes from 6 mice, one-way ANOVA, Tukey’s multiple comparisons test, the line is at the mean ± s.e.m). **e,** Total number of sheaths produced by differentiating and myelinating oligodendrocytes is similar while sheaths from myelinating oligodendrocyte are longer on average (*n* = 11 differentiating and 9 myelinating oligodendrocytes from 6 mice, unpaired t test, the line is at the mean ± s.e.m).

To formally distinguish the differentiation stages, we used a combination of morphology and Spectral Confocal Reflectance microscopy (SCoRe), a technique that enables label-free visualization of reflective, compact, myelin sheaths^30,31^. OPCs were identified as having no cellular processes that could be identified as sheaths and thus did not associate with any SCoRe signal (Fig. 2b). To distinguish between differentiating and myelinating oligodendrocytes, we quantified the SCoRe to tdTomato ratio for a subset of sheaths attached to identified cells (Fig. 2b,c; *n* = 20 cells from 6 mice). This analysis revealed a clear separation of the two groups with the newly formed sheaths from the differentiating oligodendrocytes showing little to no SCoRe signal (range 0 to 8.8%) and the sheaths from the myelinating oligodendrocytes having a more complete SCoRe coverage (range 47.2 to 93.3%) that scaled with their compaction, and therefore maturation. We further characterized the cells from these three stages with respect to cell volume, as well as process length (+ sheaths when present), and density. The differentiating oligodendrocytes showed the highest total cell and soma volume while the total process length (+ sheaths when present) was comparable between all three stages (Fig. 2d). The OPCs, in turn, had the greatest total number of branches (Fig. 2d; *n* = 10 OPCs, 11 differentiating and 9 myelinating oligodendrocytes from 6 mice, one-way ANOVA, Tukey’s multiple comparisons test). Although the differentiating and myelinating oligodendrocytes had, on average, a similar number of sheaths, the average length of the sheaths coming from myelinating oligodendrocytes was greater (Fig. 2e; *n* = 11 differentiating and 9 myelinating oligodendrocytes from 6 mice, unpaired t test).

### Mitochondrial morphology and subcellular localization differ across the oligodendrocyte lineage

To determine mitochondrial location, content, and shape across the oligodendrocyte lineage we created 3-dimensional segmentations of single cells and their mitochondria (Fig. 3-4). We identified the different cellular compartments: soma, processes (primary, secondary, tertiary+), and myelin sheaths along with their mitochondria. We then quantified the following at each stage: 1) mitochondria occupancy determined as the ratio of the total volume of mitochondria over the total volume of the whole cell or the cell compartment they occupy, 2) mitochondria density determined as the number of mitochondria per 100 µm, 3) total mitochondria number, and 4) average mitochondria length.

**Figure 3:**
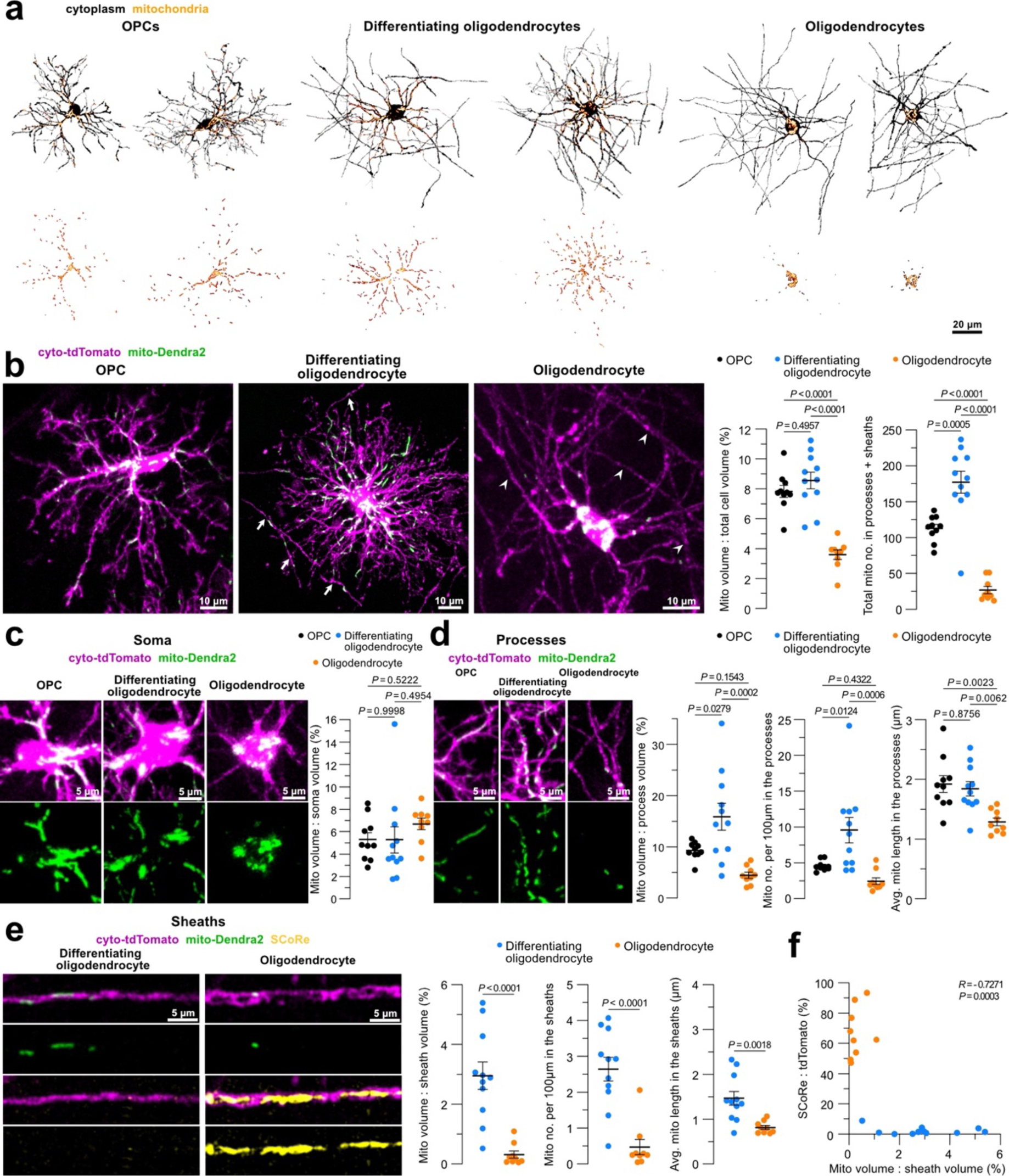
Mitochondrial morphometrics and subcellular partitioning across the oligodendrocyte lineage. **a,** 3-D reconstructions of oligodendrocyte lineage cells and their mitochondria. **b,** In vivo images of oligodendrocyte lineage cells and their mitochondria (arrows indicate non-compact sheaths; arrowheads indicate compact sheaths). Total mitochondria occupancy per cell, shown as a ratio of total mitochondria volume to total cytoplasm volume, is lowest in the myelinating oligodendrocytes. The total number of mitochondria in the processes (+ sheaths when present) is highest in the differentiating oligodendrocytes. **c,** Mitochondria in the soma of oligodendrocyte lineage cells showing no significant difference among the three stages due to variability in values within the cell groups. **d,** Mitochondria in the processes of oligodendrocyte lineage cells showing that occupancy and density is highest in the processes of differentiating oligodendrocytes, and the average mitochondria length is the lowest in the processes of myelinating oligodendrocytes (for (**b), (c), (d)**: *n* = 10 OPCs, 11 differentiating and 9 myelinating oligodendrocytes from 6 mice, one-way ANOVA, Tukey’s multiple comparisons test, the line is at the mean ± s.e.m). **e,** Mitochondria in a non-compact differentiating sheath and a compact SCoRe covered sheath from a myelinating oligodendrocyte. Sheaths from the differentiating oligodendrocytes have greater mitochondria occupancy, density, and length (*n* = 11 differentiating and 9 myelinating oligodendrocytes from 6 mice, unpaired t test, the line is at the mean ± s.e.m). **f,** Negative correlation between mitochondria distribution in myelin sheaths and myelin compaction determined by SCoRe coverage (*n* = 20 cells from 6 mice, Pearson correlation coefficient).

**Figure 4:**
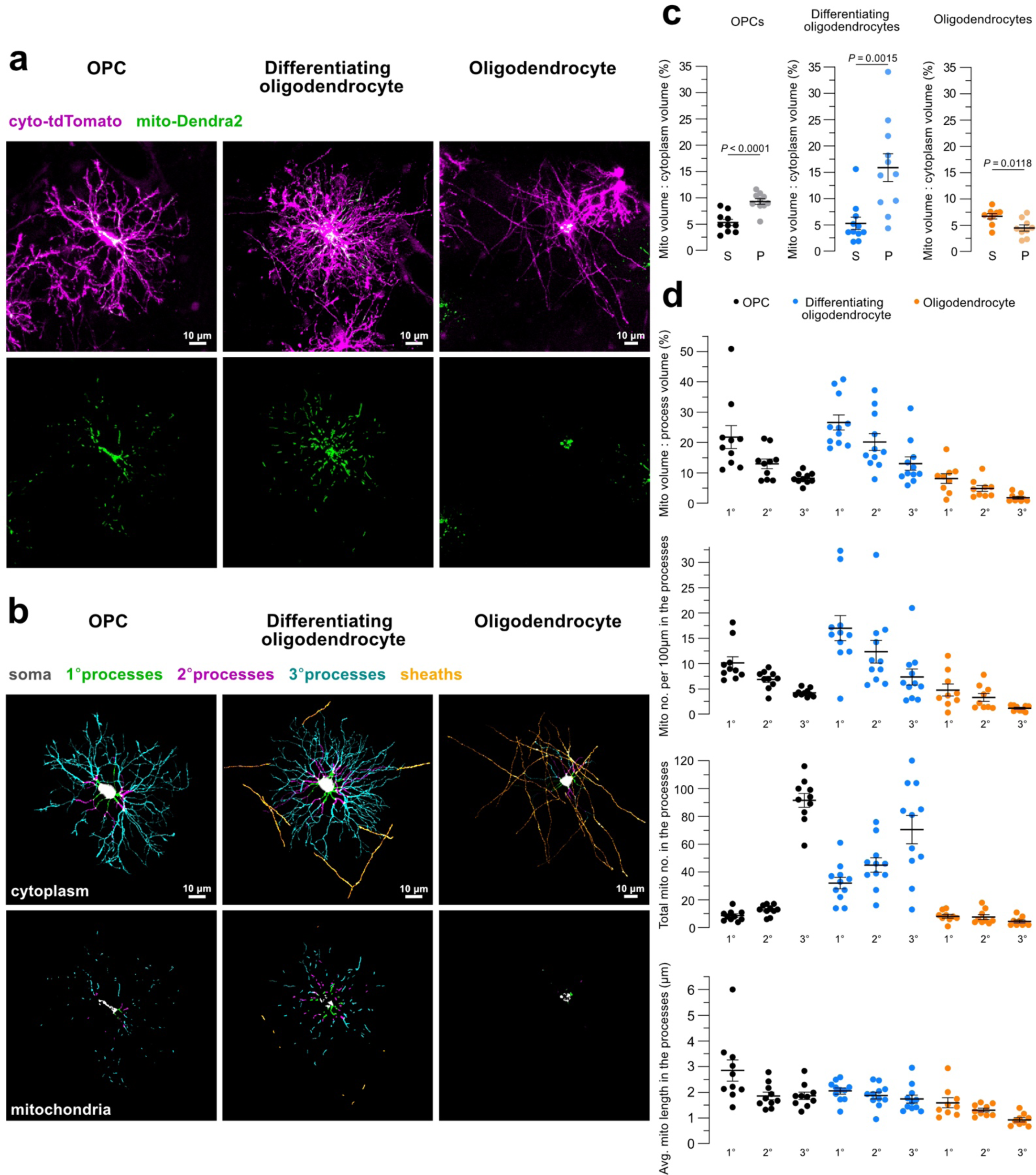
Mitochondria cellular partitioning across the oligodendrocyte lineage. **a,** Representative in vivo images of mitochondria in an OPC, differentiating oligodendrocyte, and myelinating oligodendrocyte. **b,** 3-D reconstruction of cytoplasm and mitochondria for each subcellular compartment of the cells in (**a**). **c,** Mitochondria content is higher in the processes of OPCs and differentiating oligodendrocytes compared to soma mitochondria content. The opposite is true for the myelinating oligodendrocytes (S: soma, P: processes, *n* = 10 OPCs, 11 differentiating and 9 myelinating oligodendrocytes from 6 mice, unpaired t test, the line is at the mean ± s.e.m). **d,** Quantification of mitochondria occupancy, density, total number, and average length in the primary (1°), secondary (2°), and tertiary+ (3°) processes (*n* = 10 OPCs, 11 differentiating and 9 myelinating oligodendrocytes from 6 mice, the line is at the mean ± s.e.m).

Total mitochondria occupancy was the lowest in the myelinating oligodendrocytes while the differentiating oligodendrocytes had the highest total number of mitochondria per cell (excluding soma due to difficulties in distinguishing individual mitochondria networks) (Fig. 3b). The mitochondria occupancy in the soma, however, was not significantly different between the three stages at P45 although a wide distribution of values was seen for the group of differentiating oligodendrocytes (Fig. 3c). Moreover, additional measurements of soma size and mitochondrial occupancy in OPCs and oligodendrocytes at P60 (35 days post Cre recombination) revealed significant differences between the two cell types (Fig. 5).

**Figure 5:**
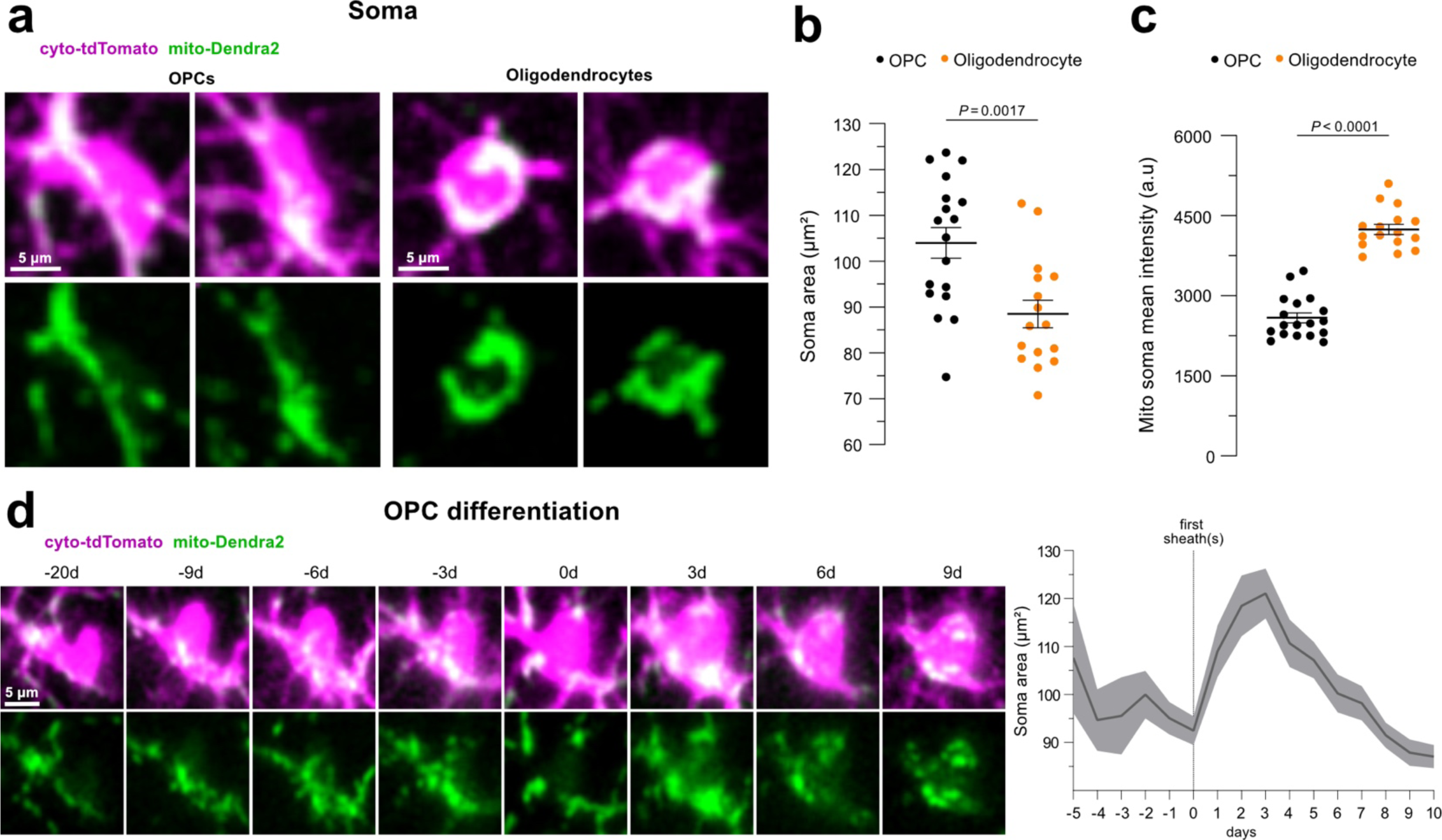
Changes in soma size and mitochondria content with oligodendrocyte maturation. **a,** Representative in vivo images of OPC and oligodendrocyte soma and their mitochondria at P60. **b,** OPC soma areas are larger. **c,** Mitochondria intensities in the somas are greater in oligodendrocytes compared to OPCs (for **(b)** and **(c)**: *n* = 18 OPCs from 6 mice and 16 oligodendrocytes from 5 mice, unpaired t test, the line is at the mean ± s.e.m, a.u: arbitrary units). **d,** Image series depicting soma changes in a differentiating OPC. Quantification of soma area over time showing an increase in soma size after sheath formation followed by a progressive decline with oligodendrocyte maturation (*n* = 7 differentiating OPCs from 4 mice, the trace represents the mean and the error bands the s.e.m).

The processes of differentiating oligodendrocytes had the highest occupancy and density of mitochondria (Fig. 3d and Fig. 4d). The shape of mitochondria in the processes changed from more elongated in OPCs to fragmented in the myelinating oligodendrocytes (Fig. 3d and Fig. 4d; for Fig. 3b,c,d: *n* = 10 OPCs, 11 differentiating and 9 myelinating oligodendrocytes from 6 mice, one-way ANOVA, Tukey’s multiple comparisons test). Lastly, the sheaths of the differentiating oligodendrocytes also exhibited higher mitochondria occupancy, density, and length compared to compact sheaths from myelinating oligodendrocytes (Fig. 3e; *n* = 11 differentiating and 9 myelinating oligodendrocytes from 6 mice, unpaired t test). Correlating the mitochondria occupancy in the sheaths to the SCoRe coverage of the cells where these sheaths originated from revealed that sheath compaction corresponds to decreased mitochondria content (Fig. 3f; *n* = 20 cells from 6 mice, Pearson correlation coefficient).

### Mitochondria morphometrics differ in human OPCs and oligodendrocytes

Next, we determined if the mitochondrial features that we observed in the mouse cortical oligodendrocyte lineage were also present in human tissue. We used a publicly available electron microscopy dataset (H01)^32^ to segment the mitochondria, cytoplasm, and nuclei in a subset of OPCs and oligodendrocytes from the temporal lobe of a human cerebral cortex sample. OPCs were identified based on their distinct process morphology, elongated shape of their soma and nuclei, and less heterochromatin compared to microglia^33^ (Fig. 6). Oligodendrocytes were identified based on their processes terminating on myelin sheaths along with their distinct soma and nuclear morphology (Fig. 6).

**Figure 6:**
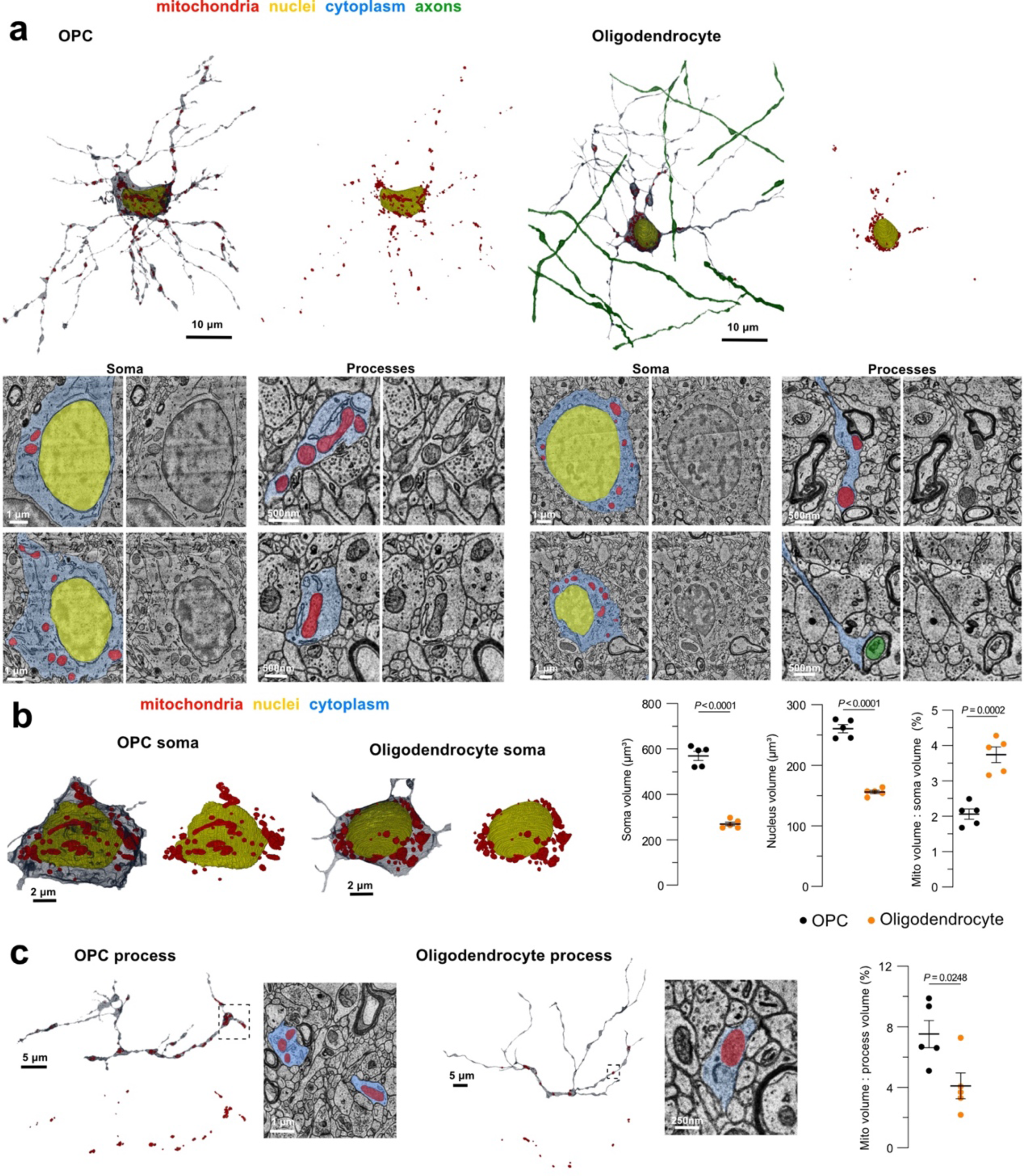
Mitochondrial morphometrics in human OPCs and oligodendrocytes. **a,** 3-D renderings, and example electron micrographs of an OPC and an oligodendrocyte captured from the temporal lobe of a 45-year-old individual showing the mitochondrial distribution and shape in the soma and processes. Several myelinated axons were reconstructed to provide context for the oligodendrocyte processes innervating the myelin sheaths surrounding these axons. **b,** Example renderings of OPC and oligodendrocyte somas, quantification of soma and nucleus volume, and mitochondrial occupancy in the soma in the two cell types (*n* = 5 OPCs and 5 oligodendrocytes, unpaired t test, the line is at the mean ± s.e.m). **c,** Example renderings of OPC and oligodendrocyte processes and quantification of mitochondrial occupancy in the processes in the two cell types (*n* = 5 OPCs and 5 oligodendrocytes, unpaired t test, the line is at the mean ± s.e.m).

Human oligodendrocyte soma volumes were smaller compared to OPCs, in part resulting in a higher soma mitochondria content in myelinating oligodendrocytes (Fig. 6b; *n* = 5 OPCs and 5 oligodendrocytes, unpaired t test). Distinct variability was observed from one cell process to the next in both cell types with some processes containing many mitochondria and others completely devoid (Fig. 6). Even with this variability, overall mitochondrial occupancy was greater in the OPC processes compared to the oligodendrocyte processes (Fig. 6c; *n* = 5 OPCs and 5 oligodendrocytes, unpaired t test). These data confirm the differences in mitochondria localization across the lineage in both the mouse and human, with preferential localization in the soma of the oligodendrocytes and in the processes of the OPCs (Fig. 3-6).

### Transient mitochondrial network reorganization during oligodendrocyte generation

After determining the mitochondria localization, content, and shape at single-time-points in different cell stages of the oligodendrocyte lineage, we next performed longitudinal imaging for 20 to 40 consecutive days at P70-P110 to track the mitochondrial changes in the same cell over time. During this period, some of the labeled OPCs spontaneously differentiated into myelinating oligodendrocytes (Fig. 7, Fig. 9b and Supplementary Video 1). We denoted the day when the first sheath/s emerged from the newly differentiated cell as day 0 in our imaging datasets. We observed that sheath compaction did not occur until ∼3 days later as demonstrated by the appearance of the SCoRe signal in single sheaths (Fig. 7b,f).

**Figure 7:**
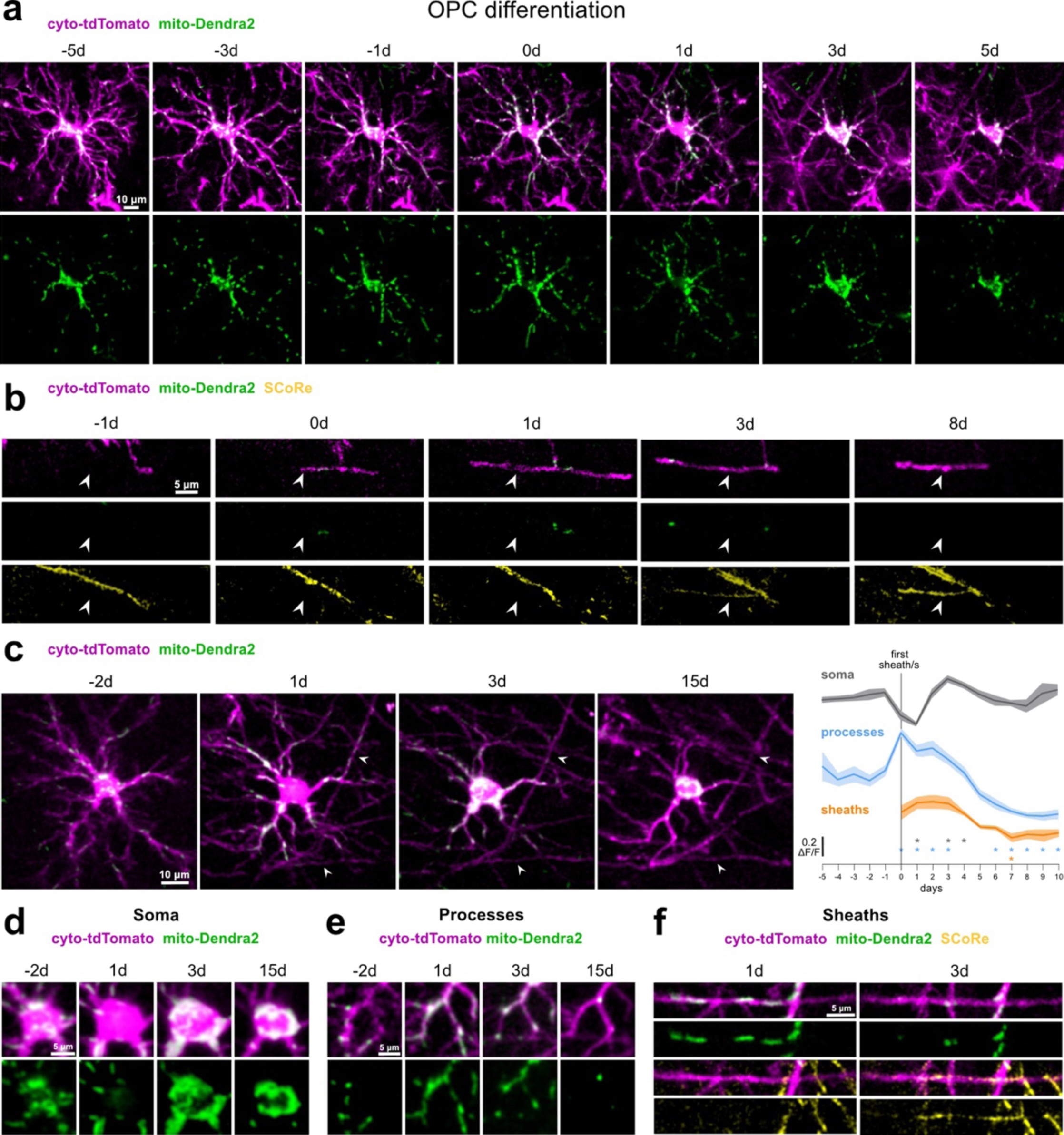
Mitochondrial network reorganization and transient expansion during oligodendrocyte generation. **a,** Longitudinal in vivo images of a single differentiating OPC and its mitochondria. Day 0 represents the first day when sheath/s are seen associated to this cell. **b,** Image series depicting a myelin sheath (arrowheads), its mitochondria and SCoRe coverage over time. The sheath forms at day 0 and compaction starts 3 days after. **c,** Representative images of a differentiating OPC and quantification of changes in mean mitochondrial intensity over 16 consecutive days (5 days before, day of and 10 days after sheath/s formation, *n* = 7 differentiating OPCs from 4 mice, the asterisks indicate data points that significantly deviate from 0 at the 99% confidence interval, the traces represent the mean and the error bands the s.e.m). Mitochondria intensity decreases in the soma coincident with an increase in the processes as sheaths are formed. 2 days later mitochondria intensity in the soma increases coincident with a drop of mitochondria in the processes and sheaths. **d, e, f,** Changes in mitochondria distribution over time in the soma **(d)**, processes **(e)**, and sheaths **(f)** of the cell in **(c)**.

To determine if there was a stereotyped progression for mitochondrial dynamics during differentiation, we quantified changes over time in mitochondrial intensity in each cell compartment (soma, processes, sheaths) for day 0, as well as 5 days before and 10 days after (Fig. 7c; *n* = 7 differentiating OPCs from 4 mice). Mitochondria intensity in the soma reached the lowest and the highest levels respectively 1 and 3 days after sheath/s emergence (Fig. 7c,d). Additionally, the mitochondrial labeling in the processes peaked on the day of sheath/s formation and decreased over time (Fig. 7c,e). Likewise, mitochondria content initially peaked in the newly formed sheaths and declined in the days following their emergence (Fig. 7b,c,f). Overall, these data show mitochondrial network reorganization and localization away from the cell soma concurrent with migration towards the newly forming sheaths. This transient state is followed by rapid disappearance of mitochondria in the distal processes and sheaths and reestablished residency in the soma during oligodendrocyte maturation and myelin compaction.

### Mitochondrial subcellular distributions are stable in non-differentiating OPCs and oligodendrocytes

To determine if there were also changes in subcellular mitochondrial localization over time for OPCs that did not change their fate for at least 20 consecutive days, OPCs that spontaneously died, or myelinating oligodendrocytes, we measured mitochondrial intensity over 10 days in the cell soma, processes, and myelin sheaths (when present) at each stage.

For the cells that remained OPCs, no significant differences were seen for the mitochondrial intensity over time in the soma and processes (Fig. 8a; *n* = 7 OPCs from 3 mice, Fig. 9a and Supplementary Video 2). Similarly, mitochondrial intensity in dying OPCs only significantly dropped the day when apoptotic bodies were detected denoted as day 10, although a rising trend was seen for the mitochondria content in the processes of the OPCs approaching death (Fig. 8b; *n* = 7 OPCs from 4 mice, Fig. 9c and Supplementary Video 3). Finally, mitochondria intensity in the soma, processes, and myelin sheaths of myelinating oligodendrocytes also remained stable over time (Fig. 8c; *n* = 7 oligodendrocytes from 4 mice).

**Figure 8:**
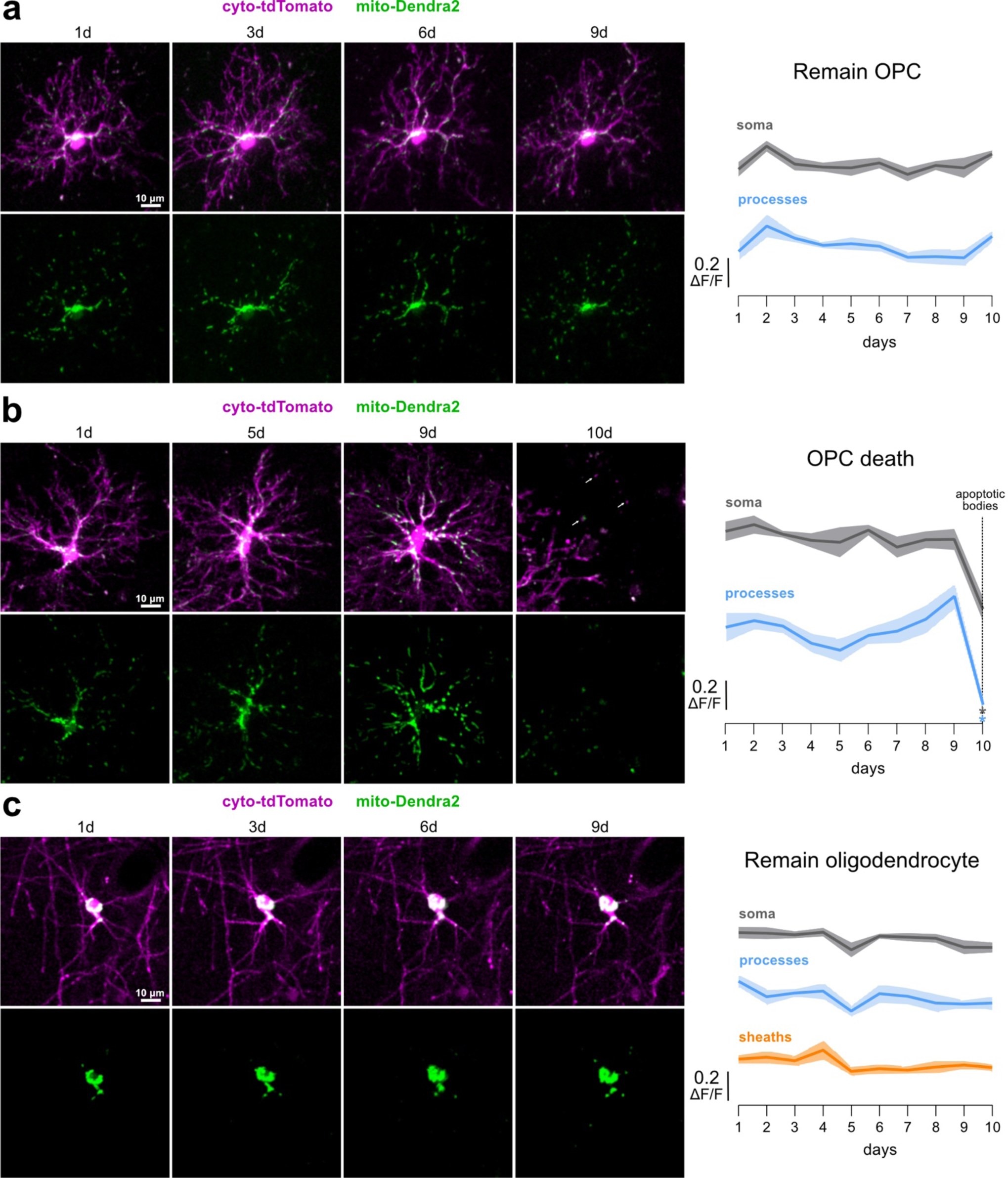
Mitochondria distribution is stable in non-differentiating OPCs and myelinating oligodendrocytes. **a,** Longitudinal in vivo images of an OPC remaining an OPC. Quantification of changes in mean mitochondrial intensity showing persistent soma and process mitochondria signal over the course of 10 days (*n* = 7 OPCs from 3 mice). **b,** Longitudinal in vivo images of an OPC that undergoes cell death where day 10 represents the day when apoptotic bodies were detected (arrows). Mitochondrial intensity 10 days prior to cell clearance showing an increasing trend of mitochondria signal in the processes before formation of apoptotic bodies and a significant drop of soma and processes mitochondria signal on day 10 (*n* = 7 OPCs from 4 mice). **c,** Longitudinal in vivo images of a myelinating oligodendrocyte. Mitochondrial intensity measurements show that mitochondrial signal remains stable in the soma, processes, and myelin sheaths of myelinating oligodendrocytes over the course of 10 days (*n* = 7 oligodendrocytes from 4 mice; for **(a), (b), (c),** the asterisks indicate data points that significantly deviate from 0 at the 99% confidence interval, the traces represent the mean and the error bands the s.e.m).

**Figure 9:**
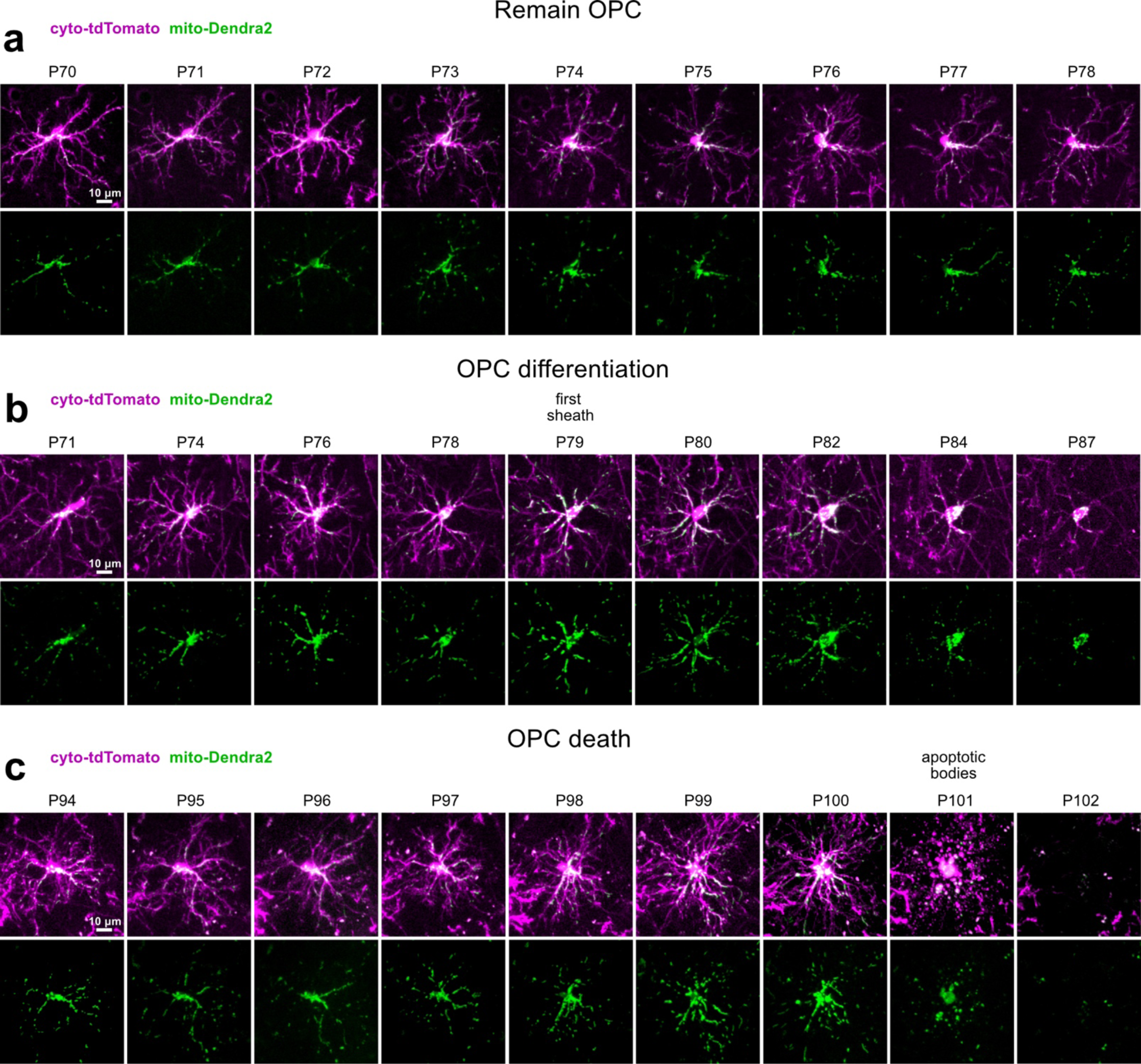
Longitudinal mitochondria dynamics across distinct OPC fates. **a,** Image series depicting an OPC that stays an OPC. Note the dynamic, yet constant distribution of mitochondria in the cell over 9 consecutive days. **b**, Image series depicting an OPC that differentiates. Note the increase in mitochondria content in the periphery as the sheaths are being made (P79) and the cell is undergoing differentiation and the subsequent decline in mitochondrial localization to the processes as the cell matures. **c**, Image series depicting an OPC that undergoes cell death. Note that mitochondria dynamics increase as the cell approaches the day of apoptosis (P101).

### Mitochondrial motility is greater in OPCs than oligodendrocytes

Next, we tracked the mitochondrial dynamics in the OPCs and oligodendrocytes over hours to minutes. To first determine general mitochondrial movement over hours, we photoconverted the mito-Dendra2 fluorescence in the soma of OPCs or oligodendrocytes and followed these cells over time at 4, 24, and 48 hours (Fig. 10). The intensity of the photoconverted mitochondria dropped quickly at 4 hours for the OPCs but remained stable for the oligodendrocytes. Similarly, the decline was greater for the OPCs than for the oligodendrocytes at 24 and 48 hours (Fig. 10c,d; *n* = 11 OPCs and 12 oligodendrocytes from 4 mice, multiple unpaired t tests with Holm-Šídák method). This higher rate of dilution of the photoconverted mitochondria signal in the OPCs suggests higher dynamics and mitochondrial motility in these cells.

**Figure 10:**
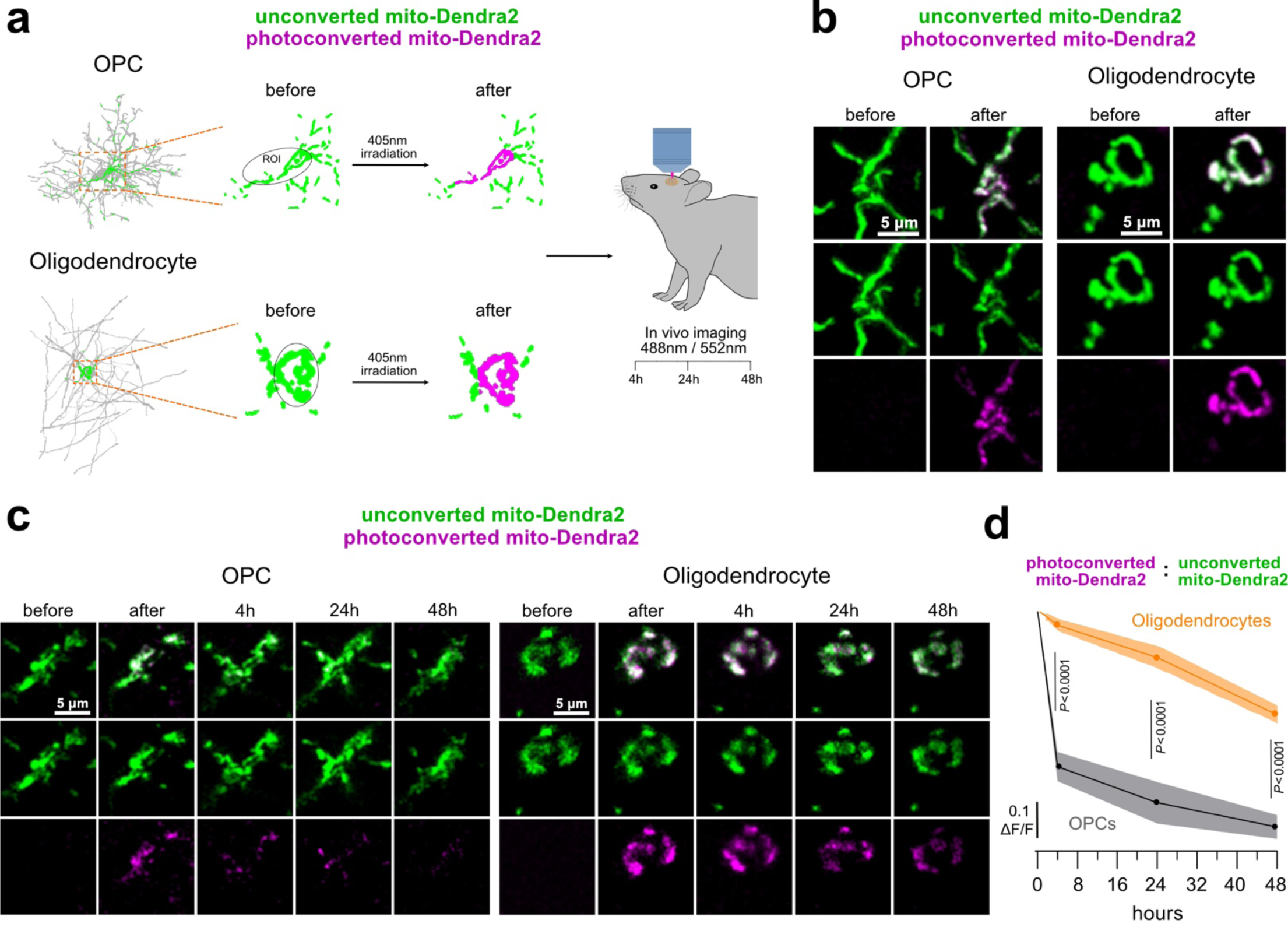
Mitochondrial network stability differs between OPCs and oligodendrocytes. **a,** Schematic illustration of photoconversion experiments. A circular ROI was drawn around the soma of mito-Dendra2-only labeled OPCs and myelinating oligodendrocytes and irradiated with a 405 nm laser. Unconverted and photoconverted mito-Dendra2 signal was imaged in vivo before, immediately after, and at 4, 24 and 48 hours after photoconversion. **b,** Example images of mitochondria signal in an OPC and an oligodendrocyte before and after photoconversion. **c,** Longitudinal images of mitochondria before, after and at 4, 24, and 48 hours after photoconverting soma mitochondria in an OPC and an oligodendrocyte. **d,** Normalized photoconverted mitochondria ratios within the soma ROI at 4, 24 and 48 hours after photoconversion showing a rapid loss in photoconverted mitochondria in the OPCs while photoconverted mitochondria in the oligodendrocytes persist for days (*n* = 11 OPCs and 12 oligodendrocytes from 4 mice, multiple unpaired t tests with Holm-Šídák method, the traces represent the mean and the error bands the s.e.m).

To further test this hypothesis, we imaged cells once a minute over 30 minutes in awake mice and tracked mitochondrial motility in the OPCs and oligodendrocytes. Throughout this timeframe, we observed a mixture of motile and stationary mitochondria in the OPCs (Fig. 11a,b and Supplementary Video 4) and only stationary mitochondria in the oligodendrocytes (Fig. 11c,d and Supplementary Video 5). The average mitochondria displacement per cell was higher in the OPCs compared to oligodendrocytes (Fig. 11e). On average, 11% of mitochondria in the OPCs traveled above a displacement threshold set to eliminate measurements of mitochondrial oscillations and any awake imaging movement artifacts, while no mitochondria in the oligodendrocytes traveled above the same displacement threshold (Fig. 11e; *n* = 18 OPCs from 6 mice and 16 oligodendrocytes from 5 mice, unpaired t test).

**Figure 11:**
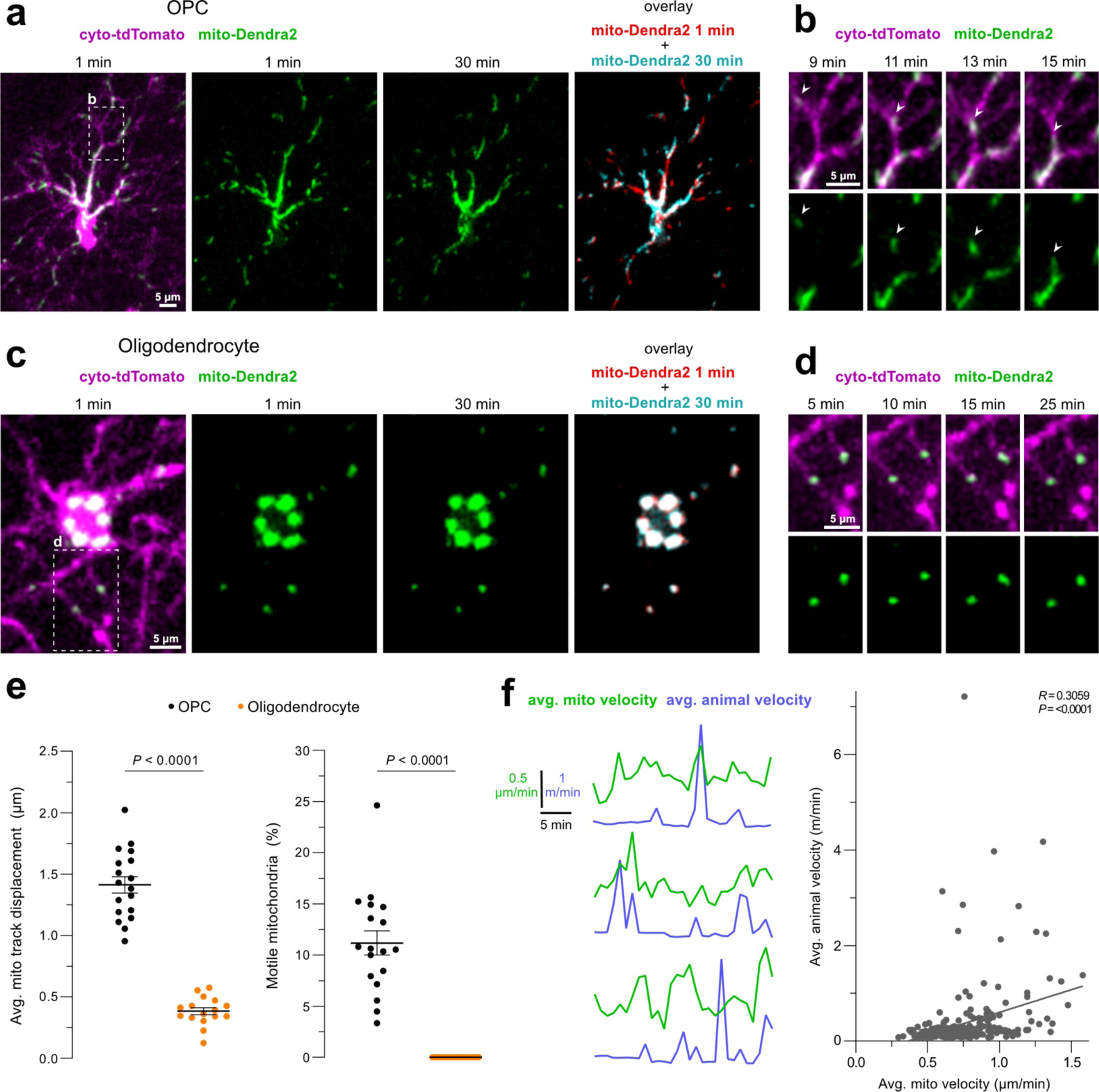
Mitochondria are motile in OPCs and stationary in oligodendrocytes. **a, c,** Representative in vivo images of an OPC **(a)** and an oligodendrocyte **(c)** at the first and last minute of a 30-minute time series captured from an awake mouse. The overlay of mitochondria signal at these two time points shows numerous motile mitochondria in the OPC but none in the oligodendrocyte. **b,** The boxed region in **(a)** between minute 9 and 15 of the time series. Arrowheads point at mitochondria moving in the OPC processes. **d,** The boxed region in **(c)** showing stationary mitochondria in the oligodendrocyte between minute 5 and 25 of the time series. **e,** Mitochondria displacement is greater in the OPCs compared to oligodendrocytes. Percentage of mitochondria per cell that displace ≥3 microns over the 30-minute time series, showing that mitochondria in OPCs are motile while oligodendrocyte mitochondria are stationary (*n* =18 OPCs from 6 mice and 16 oligodendrocytes from 5 mice, unpaired t test, the line is at the mean ± s.e.m). **f,** Representative traces of average animal velocity and average velocity of mitochondria within OPCs. Correlation between animal locomotion velocity and mitochondria velocity (*n* = 261 events captured from 31 OPCs from 9 animals across the 30-minute time series, Pearson correlation coefficient).

To understand what external factors could have influenced the range of mitochondria motility in the OPCs (Fig. 11e), we considered both animal sex and animal locomotion during the image acquisition. Animal sex was not a contributing factor revealing no differences in OPC motility between female and male mice (*n* = 19 OPCs from 6 female and 38 OPCs from 11 male mice, unpaired t test, *p* = 0.7255). On the other hand, animal locomotion itself was linked to mitochondria motility in the OPCs. Comparing the average animal locomotion velocity and mitochondria velocity per every minute of the imaging timeframe revealed a weak correlation between the two (Fig. 11f; *n* = 261 events captured from 31 OPCs from 9 animals across the 30-minute time series, Pearson correlation coefficient). These data suggested that animal locomotion was partially associated with higher OPC mitochondria motility.

### Arousal state influences OPC mitochondrial motility

With animal locomotion partly contributing to mitochondria motility in the OPCs, we next determined if animal arousal state impacted mitochondrial motility by treating animals with anesthetics or sedatives during the imaging experiments (Fig. 12a). OPC mitochondrial motility decreased under both isoflurane (Fig. 12b and Supplementary Video 6; *n* = 18 OPCs from 6 mice, paired t test) and ketamine/xylazine (Fig. 12c and Supplementary Video 7; *n* = 17 OPCs from 5 mice, paired t test) induced anesthesia.

**Figure 12:**
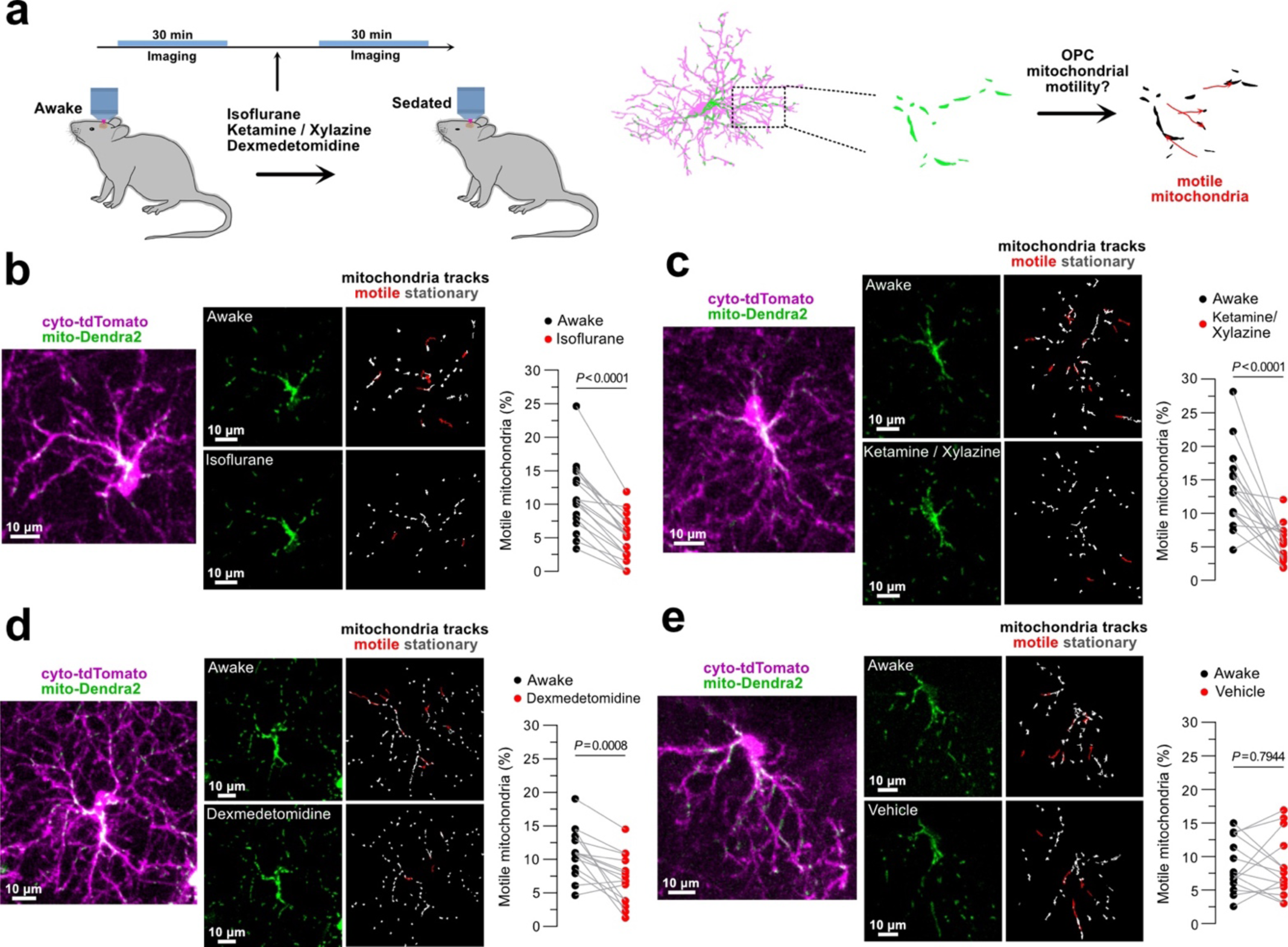
Animal sedation decreases mitochondria motility in OPCs. **a,** Schematic illustration of experimental timeline. Animals were imaged for 30 minutes in awake and anesthetized/sedated conditions and mitochondria in the OPCs were analyzed to determine their motility rate. **b, c, d, e,** Left of each panel: representative images of mitochondrial subcellular distribution in the analyzed OPCs at the start of the awake imaging session. Right of each panel: mitochondria signal at the start of each imaging session in each condition along with mitochondria tracks only in the processes of the targeted OPCs across 30 minutes for the awake and 0.8-1% isoflurane (**b**, *n* = 18 OPCs from 6 mice, paired t test), 100 mgkg^-1^ ketamine / 10 mgkg^-1^ xylazine (**c**, *n* = 17 OPCs from 5 mice, paired t test), 0.5 mgkg^-1^ dexmedetomidine hydrochloride (**d**, *n* = 15 OPCs from 4 mice, paired t test) and vehicle conditions (**e**, *n* = 14 OPCs from 4 mice, paired t test). Tracks in red represent mitochondrial displacement of 3 µm or greater and are referred to as “motile”. All the conditions in **(b)**, **(c)**, and **(d)** reduced mitochondria motility compared to awake conditions.

Norepinephrine is one of the key neuromodulating signals that change with locomotion and under anesthesia, thus, to test the effects of sedation in addition to anesthesia we next blocked norepinephrine release with dexmedetomidine hydrochloride. Consistent with the anesthetics, dexmedetomidine treatment decreased OPC mitochondrial motility (Fig. 12d and Supplementary Video 8; *n* = 15 OPCs from 4 mice, paired t test) while no change was observed in vehicle-treated animals (Fig. 12e and Supplementary Video 9; *n* = 14 OPCs from 4 mice, paired t test). These results reveal that the animal arousal state influences OPC mitochondria motility and opens a new target for manipulating mitochondrial dynamics in these cells.

### Aged OPCs display reduced mitochondrial length and motility

Given the importance of mitochondria for cellular homeostasis and differentiation, we next characterized the mitochondrial morphology and distribution in aged OPCs which have a reduced capacity to differentiate. We segmented in 3-D a subset of young and aged OPC processes and their mitochondria from high resolution single-time-point images coming from mice aged either 1 month or 18-21 months (Fig. 13a). Sholl analyses revealed fewer intersections in aged OPCs (Fig. 13b). The mitochondrial volume occupancy was decreased in aged OPCs while the density did not change (Fig. 13b). On the other hand, aged OPCs displayed shorter mitochondrial length, likely accounting for the reduced mitochondria volume in these cells (Fig. 13b; *n* = 18 OPCs from 4 young mice and 18 OPCs from 6 aged mice, unpaired t test). Finally, we imaged mitochondria motility for 30 minutes in awake aged mice. While the percent mitochondria moving in young and aged OPCs did not reach a statistically significant difference between the two groups, aged OPC mitochondria exhibited a lower average displacement suggesting decreased overall mitochondrial motility within the aged OPCs (Fig. 13c; *n* = 18 OPCs from 6 young mice and 22 OPCs from 3 aged mice, unpaired t test and Supplementary Video 10). Thus, aged OPCs contained shorter mitochondria that moved less.

**Figure 13:**
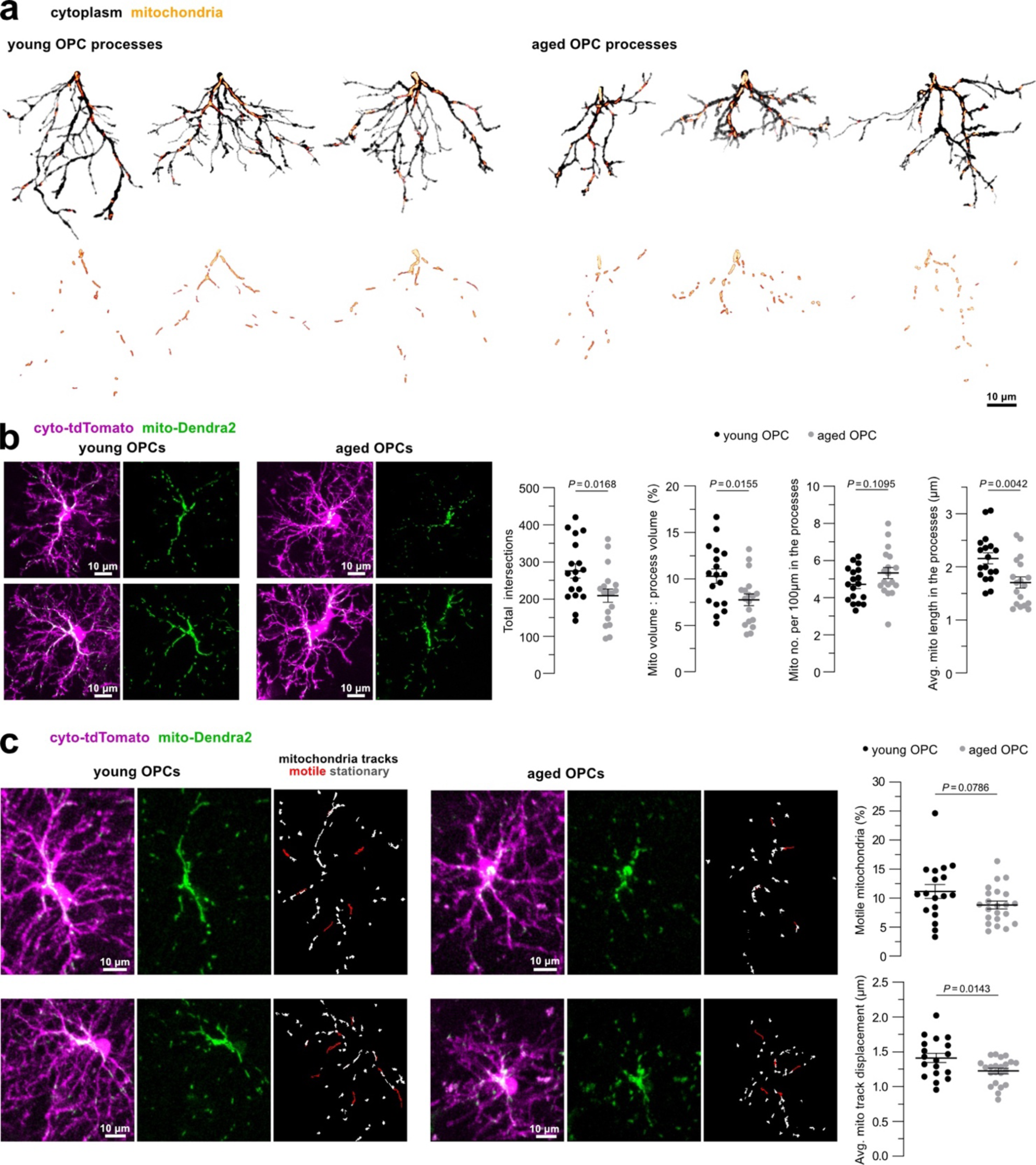
Aged OPCs have shorter mitochondria that are less motile. **a,** 3-D reconstructions of a subset of processes and their mitochondria in young and aged OPCs. **b,** In vivo images of young and aged OPCs and their mitochondria. Aged OPCs display a lower number of total intersections as quantified by Sholl analysis. Mitochondria volume fraction is also lower in the processes of aged OPCs while mitochondria density shown as number of mitochondria per 100 µm is not statistically different. Mitochondria in the processes of aged OPCs are shorter (*n* = 18 OPCs from 4 young mice and 18 OPCs from 6 aged mice, unpaired t test, the line is at the mean ± s.e.m). For **(a)** and **(b)**, young and aged OPCs refer to OPCs coming respectively from mice aged 1 month and 18-21 months. **c,** Representative images of mitochondrial subcellular distribution in OPCs at the start of the awake imaging session along with mitochondrial tracks in the analyzed OPC processes that show mitochondria displacement along the 30-minute imaging series. Tracks in red represent mitochondrial displacement of 3 µm or greater and are referred to as “motile”. The percentage of mitochondria that move at or above a 3 µm threshold and the average displacement of mitochondria in the processes of the aged OPCs (*n* = 18 OPCs from 6 young mice and 22 OPCs from 3 aged mice, unpaired t test, the line is at the mean ± s.e.m). Young and aged OPCs refer to OPCs coming respectively from mice aged 2 months and 18-21 months.

## DISCUSSION

The organelle dynamics that take place before, during, and after fate transitions in the oligodendrocyte lineage are largely unknown. In this study, we generated a transgenic mouse that enables visualization of mitochondria specifically in the oligodendrocyte lineage. Using high-resolution fluorescence optical imaging coupled with intravital longitudinal and label-free approaches, we characterized how mitochondria content, shape, and motility change in different stages and fates of the oligodendrocyte lineage. During OPC differentiation, mitochondria content in the processes increased as sheaths were being made and dropped as sheaths formed compact myelin. This mitochondrial redistribution was specific to differentiation as mitochondria were stable during other OPC fates. Mitochondria shape transitioned from more elongated in OPCs to fragmented in myelinating oligodendrocytes. In addition, oligodendrocyte mitochondria were stationary while OPC mitochondria were more dynamic. Animal sedation decreased OPC mitochondria motility and aged OPCs had shorter mitochondria that moved less. Overall, these data reveal that mitochondria are dynamically positioned within OPCs undergoing fate decisions in the live cerebral cortex. This provides new insight into identifying important temporal windows for when these changes occur and how subcellular organelle dynamics could impact oligodendrocyte generation.

Mitochondria distribution, shape, and motility are closely linked to mitochondria function^10,34–37^. The increase in mitochondria content during OPC differentiation can be attributed to the role of mitochondria in providing energy and substrates for lipid synthesis^38^. Indeed, high levels of ATP are needed for the formation of myelin sheaths^39,40^ and impairing mitochondria has been shown to reduce myelin lipids in the PNS^41^. Given the long life of myelin components^1,42^, the energy requirements for maintaining the myelin sheaths are likely lower than for synthesizing them which could explain the drop in mitochondrial content as sheaths mature. As myelin sheaths compact, they exclude the cytoplasmic content and hence reduce the available space for the organelles which is consistent with the decline in mitochondria content with compaction. Like mitochondria, peroxisomes have also been described in regions of non-compact myelin^43^.

At our single-time-point analysis, we were not able to distinguish exactly when changes in mitochondria content took place as we labeled and analyzed a continuum of cell states in the oligodendrocyte lineage, thus explaining some of the data variability particularly in the differentiating oligodendrocytes group (Fig. 3). Longitudinal imaging clarified when and how mitochondrial partitioning occurred and confirmed that even within days mitochondria content can change drastically during oligodendrocyte generation (Fig. 7). For example, we saw that mitochondria in the soma of the differentiating oligodendrocytes reached both the lowest and highest mitochondrial content within 3 days. A potential reason for mitochondria leaving the soma is to assist in the tips of the processes as the sheaths are formed and initiate compaction. In support of this, an earlier study suggested that mitochondria in the processes are more active at this stage than the mitochondria in the soma^44^. Once the sheaths are formed and compact, mitochondria either return to the soma and/or get locally degraded which, due to technical limitations, we have not been able to distinguish. Mitochondria degradation could explain the decline in mitochondrial content during oligodendrocyte differentiation which occurs over several days until it reaches a low steady state density. Consistent with this, previous work suggests that mitophagy is a prerequisite for oligodendrocyte generation^22^. Moreover, mitochondrial fragmentation has been shown to precede and facilitate mitophagy^45^ thus the maturation-dependent decline in mitochondrial length is also consistent with mitophagy.

The ultrastructural analyses in the human tissue revealed differences between OPCs and oligodendrocytes with regards to soma and nucleus volume, and mitochondrial subcellular partitioning (Fig. 6). Initially these measurements appeared inconsistent with the single-time-point mouse data showing trends towards differences between these cell populations but not reaching significance (Fig. 2-3). Importantly, these measurements from the mouse were taken from cells 20 days after Cre recombination (tamoxifen at P25 and imaging at P45) meaning the myelinating oligodendrocytes labeled and analyzed were not more than ∼15 days “old”, with many likely to have differentiated less than 10 days prior to the analyses. While these cells had canonical myelinating oligodendrocyte morphology and were making compact myelin, they were still relatively young, particularly when comparing the age of the cells in the 45-year-old human tissue. Subsequent longitudinal analysis in the mouse revealed changes in soma size during in vivo OPC differentiation with an initial peak immediately following the formation of the first sheath/s followed by a progressive drop as the cell matured (Fig. 5). Moreover, additional measurements of soma size and mitochondrial occupancy in OPCs and oligodendrocytes at P60 (35 days post Cre recombination) revealed significant differences between the two cell types consistent with the human data (Fig. 5). Overall, these observations reveal a stereotyped and progressive cell morphology change and redistribution of mitochondria within the differentiating cell over weeks. There are abrupt (within hours and days) changes occurring throughout the differentiation event and initial myelin sheath formation, but mitochondrial partitioning continues to gradually change for weeks after oligodendrocyte generation.

Changes in mitochondria content and shape during oligodendrocyte differentiation are likely interdependent with metabolic transitioning that occurs between high rates of oxidative phosphorylation in OPCs^46,47^, increased oxidative metabolism and higher ATP levels in differentiating oligodendrocytes^29^, and a switch to glycolysis in oligodendrocytes^48,49^. Likewise, longer mitochondria are generally associated with increased rates of oxidative phosphorylation^50,51^ although this varies depending on the cell type and cellular context^34,52^. In addition to providing metabolic substrates, mitochondria could also be buffering calcium in defined subcellular compartments. Mitochondria localized in the processes of the OPCs and immature oligodendrocytes are thought to be important for maintaining calcium homeostasis^44,53^. In addition, mitochondria have been linked to calcium transients in newly forming myelin sheaths which decline with myelin maturation^54^. Lastly, mitochondria are a central player of cellular death. High mitochondria content during oligodendrocyte differentiation could be linked to increased susceptibility towards oxidative stress and mitochondria disruption at this stage^19,29,55^. Our longitudinal imaging showed a trend toward increased mitochondria content in the processes of the OPCs before the formation of apoptotic bodies and death. While not statistically significant based on our cutoff (Fig. 8b), it is likely that this trend signifies the initiation of an oligodendrocyte differentiation program but a failure to survive and integrate due to unfavorable intra- and extracellular signals.

Mitochondria move within the cell to relocate to the sites where they are needed^35–37^. Our data show that mitochondria motility and overall dynamics were higher in OPCs compared to myelinating oligodendrocytes (Fig. 10-11). Consistent with this, mitochondria motility in neurons decreases with their maturation^56,57^ and corresponds to increased levels of syntaphilin^58^, a mitochondrial docking protein whose levels also change with oligodendrocyte maturation^59^. Moreover, mitochondria motility in neurons and astrocytes in vivo is lower compared to in vitro or ex vivo slice studies^56,60–62^. Similarly, we saw that oligodendrocyte mitochondria were more stationary in vivo than what had been reported in slices^25^ and cultured oligodendrocytes^26^. Other than the physiological context, the maturation state of these cells could also contribute to this discrepancy as the age of the oligodendrocyte, even if expressing many mature oligodendrocyte markers (as discussed above), is likely different between these studies. Additionally, the in vivo system better models the neuronal activity and neurotransmitter release that happens in the native environment of these cells. Our results suggest that arousal state plays a role in modulating mitochondria motility in OPCs (Fig. 12). Arousal state has been linked to changes in calcium transients in OPCs and other glia^62–66^. Thus, arousal-dependent changes in mitochondria motility is consistent with recent evidence of decreased calcium dynamics in OPCs in anesthesia conditions or when norepinephrine release is blocked^63^. Norepinephrine can influence the differentiation and/or proliferation of oligodendrocytes and the expression of norepinephrine receptors in the OPCs changes with their differentiation^63,64,67^. Overall, these observations suggest a potential connection between mitochondria and calcium dynamics and raise questions for how animal arousal state influences mitochondria in these cells and therefore the cell fate in the oligodendrocyte lineage.

Consistent with reduced mitochondrial trafficking with aging in other cells^68^, including neurons^69^, aged OPC mitochondria also displayed reduced displacement. In addition, the reduced mitochondrial volume fraction and size could explain the decline in ATP levels and cellular respiration observed in cultured OPCs from aged animals^29^. Aging is associated with increased oxidative stress and decreased mitophagy which could lead to mitochondrial fragmentation and accumulation of damaged mitochondria^27^. This can partially explain why there was a decrease in size of the individual mitochondrial networks and no changes in the mitochondrial density in the aged OPCs. It is also important to note that the mitochondrial morphometrics and motility were variable in the aged OPCs, likely reflecting the documented heterogeneity of OPC states in the aged brain^70^. Whether these are linked to functional heterogeneity between these cells is yet to be determined. Nonetheless, these data point to overall metabolic changes occurring in aged OPCs that would likely impact their ability to successfully generate new oligodendrocytes.

## Acknowledgments

We thank members of the Hill and Hoppa laboratories at Dartmouth College for valuable feedback on this work. We would also want to thank Dr. Shahroz Khan for helping with data analysis. This study was supported by National Institutes of Health grant R01NS122800 and the Esther A. & Joseph Klingenstein Fund and Simons Foundation to R.A.H. and American Heart Association grant. 23PRE1018862, https://doi.org/10.58275/AHA.23PRE1018862.pc.gr.161154 to Xh.B.

## Author contributions

Xh.B and R.A.H designed and performed all the experiments, analyzed the data, wrote the manuscript, and secured funding. R.A.H supervised the study.

## METHODS

### Animals

The following lines: *Cspg4-*creER^71^ (JAX #008538); Ai9 ^72^ (JAX #007909); PhAM ^73^ (JAX #018385) were bred to generate the *Cspg4-*creER: Ai9: PhAM triple transgenic mice. This mouse line allows for conditional visualization of mitochondria (mito-Dendra2 labeled) and cytoplasm (tdTomato labeled) in cells expressing the chondroitin sulfate proteoglycan 4 (Cspg4, also known as NG2) and their progeny. A single dose of tamoxifen (0.4-0.8 mg) was intraperitoneally injected at P25. This produced sparse dual or single recombination providing double and single tdTomato-only or mito-Dendra2-only labeled cells. Both male and female mice aged P30-P651 were used for the experiments. All animals were housed in a temperature and humidity-controlled vivarium in a 12h light/dark cycle with ad libitum access to food and water. All the animal procedures were approved by the Institutional Animal Care and Use Committee at Dartmouth College.

### Surgical procedures

In vivo imaging was performed in acute and chronic cranial windows implanted over the somatosensory cortex. First, the animals were anesthetized via intraperitoneal injections of ketamine/xylazine (100 mgkg^-1^/ 10 mgkg^-1^) or placed under 1% isoflurane. After sterilizing and removing the skin covering the skull, a craniotomy was performed over a 3 by 3 mm area of the exposed skull. The skull and the underlying dura were removed and a circular #0 glass coverslip was positioned on top of the craniotomy region to create an optically accessible imaging window^74^. Carprofen analgesic (5 mgkg^-1^) was administered subcutaneously before and after the surgery as well as in the following 24 and 48 hours. Acute imaging was performed on the day of the surgery after the animals recovered from anesthesia. Chronic imaging was performed after allowing at least 3 weeks for the animals to acclimatize to the surgery.

### Imaging

For all anesthetized in vivo imaging experiments, the animals were placed under 0.8-1% isoflurane anesthesia (2% used for induction) and imaged on an upright laser-scanning confocal (Leica SP8) or upright laser-scanning two-photon microscope (Bruker Ultima) with a 20X water immersive objective (respectively Leica NA 1.0 and Zeiss NA 1.0). The tdTomato was excited by 552 nm laser wavelength on the confocal and 1040 nm on the two-photon. Unconverted mito-Dendra2 was excited by 488 nm and 920 nm laser wavelengths from the respective microscopes. SCoRe images were captured on the confocal microscope by overlaying the reflectance from 448 nm, 488 nm, 552 nm, and 637 nm lasers^30,31^. The Z-stacks were taken over a depth of 50-100 μm in layer I of the somatosensory cortex with a 1.5µm step size. The same settings were used for the awake imaging in which the animal’s head was fixed in the Neurotar’s Mobile HomeCage setup and the animal could move ad libitum on an air table.

### 3-dimensional (3-D) reconstructions and single-time-point analysis

3-D reconstructions were used to quantify the mitochondrial density and volume fraction (total mitochondrial volume divided by the total cytoplasmic volume) for each cell and their compartments (soma, primary, secondary, tertiary+ processes, and myelin sheaths) along with mitochondria, processes, and myelin sheaths lengths. We traced a group of cells from single-time-point, high-resolution images, captured from P45-P46 animals following acute cranial window and later separated them into 3 categories: OPCs (polydendritic morphology lacking myelin sheaths), differentiating oligodendrocytes (presence of premature sheaths with less than 34.54% SCoRe coverage), mature myelinating oligodendrocytes (presence of compact myelin sheaths with more than 34.54% SCoRe coverage). For identification of premature myelin sheaths that completely lacked SCoRe signal, the following criteria were used: be a terminal process (no branching), be at least 5 µm in length, and have a larger diameter than the proximal process.

The cells of interest were cropped, smoothed, and traced in 3-D using the Simple Neurite Tracer (SNT, version 4.1.2) Fiji Plugin^75,76^. Tracing was done for all the cell compartments for which we could unambiguously determine that they were attached to the cell. The soma was marked by a single-point path placed in the center. A segment path was created along the length of the processes and myelin sheaths. A separate path was created at every branching point and the order of branching was noted. The initial processes emerging from the soma were considered as primary, their immediate branches were considered as secondary, and the branches coming from secondary processes and above were denoted as tertiary+. Only the myelin sheaths for which we could assign a myelinating process were traced. Similarly, mitochondria were traced along their length throughout every cell compartment and noted accordingly to which compartment they belonged. All the traced paths were then filled using the “Fill out” command from the SNT Path Manager menu. A threshold was set so that all the corresponding signals along the path were accurately included in the fill. Different thresholds were set to match the thickness of the traced structure. The volume values for the cytoplasmic and mitochondrial channels were exported from the fills. Additionally, the path lengths were measured using the “Measure path” command from the SNT Path Manager menu. Similarly, SNT reconstructions were generated to analyze the images captured from young and aged animals (respectively P30-P35 and P539-P651). In this case, a subset of processes was selected per cell, including a primary process emerging from the soma and all the subsequent branches. The experimenter was blinded to the age of the animal and mitochondrial labeling when selecting the processes to be analyzed. Sholl analysis was performed in Z-projections of reconstructed images using the SNT “Sholl Analysis” command^77^ to determine the total number of intersections. The tip of the primary process was set as the center, the start radius was set at 0 µm, the step size was set at 1 µm and the end radius at 70 µm.

The single-time-point soma area analysis was done for cells at P60 (5 weeks post tamoxifen administration). Z-projections were created from the cropped stacks of cells of interest and smoothed. A circular region of interest (ROI) was drawn around the cell soma using only the cytoplasmic channel as a reference. The area and mean intensity of mitochondrial signal within the ROI were measured in Fiji.

### SCoRe analysis

SCoRe coverage was analyzed to determine the level of compaction in the newly formed and mature myelin sheaths coming from differentiating or myelinating oligodendrocytes. The SCoRe channels were merged into a single channel which was later smoothed and thresholded by running the Robust Automatic Threshold Selection in Fiji. The same threshold method was applied for the cytoplasmic channel. Only a sample of the sheaths that had no overlapping signal (from either background or other sheaths and structures in the territory) were traced using SNT and filled out for both the SCoRe and the cytoplasmic channels to generate ROIs of the sheath of interest. The mean intensity within the given ROIs were measured and ratios of the SCoRe to cytoplasmic signal in the sheaths were analyzed. K-means clustering was run to determine the threshold for high versus low SCoRe coverage in our dataset (k=2, threshold = 34.54%).

### Longitudinal analysis

To observe mitochondrial dynamics occurring with cell fate changes, we performed longitudinal imaging for 20 to 40 consecutive days between P70 to P110. Out of this, data analysis was done for 16 consecutive days for OPCs that differentiate (5 days before, the day of, and 10 days after sheath formation) where day 0 was equivalent to the day when the first sheath/s emerged from the cell. For cells that did not change their fate (remained OPCs or myelinating oligodendrocytes), data analysis was done for 10 consecutive days out of at least 20 time points where these cells’ fate remained unchanged. For cells that die, analysis was done for 10 consecutive days before the cell signal disappeared where the 10^th^ day was the day that apoptotic bodies were detected.

To quantify changes in mitochondrial intensity over time, Z-projections that spanned all the parts of the cell that were to be analyzed were created at each time point. The Z-projections were combined into a hyperstack, registered, and smoothed. ROIs were drawn using the Fiji selection brush tool along the cell soma and 3 primary processes, 3 secondary processes, and 3-4 myelin sheaths (when present) per cell. The cellular compartments were selected independently of the mitochondria channel and traced over time with reference only to the cytoplasmic channel. At the end of the tracing, they were visually inspected and traced compartments that had overlapping mitochondrial signals coming from other cells in the environment were excluded from the ROI. If the same process was not found in all the time points due to cellular morphological changes, it was replaced by tracing a different emerging process of the same branching order from the same cell. The area and mean fluorescence intensity from the mitochondria label were measured within each ROI and the data were normalized to the average mean fluorescence intensity of all the data points analyzed per cell compartment.

The average data for all 6 traced processes (3 primaries, 3 secondaries) and 3-4 myelin sheaths per cell were plotted.

### Electron microscopy dataset analysis

To investigate mitochondria ultrastructure and determine if mitochondrial differences in the oligodendrocytes and their precursors were present in the human cortex, we used the H01 dataset available https://h01-release.storage.googleapis.com/data.html^32^. The authors of this dataset used a non-pathological temporal lobe sample obtained from a resection surgery of a 45-year-old epileptic patient, sectioned 5000 slices at 30-40 nm thickness, and imaged them using scanning electron microscopy^32^. They also identified the cell types and computationally segmented their structure in 3-D which can be viewed in Neuroglancer^32^. We used the Volume Annotation and Segmentation Tool (VAST Lite, version 1.4.1)^78^ to open the H01 dataset, import the coordinates of our cells of interest taken from Neuroglancer, and fill the cell cytoplasm that had already been automatically segmented. We selected a sample of 5 oligodendrocytes and 5 OPCs and re-confirmed their identity with OPCs having ramified processes, an elongated soma and nucleus, and less heterochromatin than microglia^33^ and oligodendrocytes having processes ending on myelin sheaths. We visually inspected and corrected the cytoplasmic fills as needed and analyzed the cell soma and the connected processes separately. Next, we segmented all we could identify as mitochondria in the somas and a subset of processes for each cell type (a total of 26 primary OPC processes and all their branches and a total of 14 primary oligodendrocyte processes and all their branches). We exported the cytoplasmic and mitochondria volumes from the region of interest using VAST and calculated mitochondria occupancy as the ratio of total mitochondria volume to total cell compartment volume.

### Photoconversion

Photoconversion was done in mito-Dendra2-only labeled cells of 2–3-month-old animals to characterize changes in mitochondrial dynamics in OPCs and myelinating oligodendrocytes. We referred to changes in mitochondrial shape and content to determine the state of the cell (OPCs: higher content of mixed shape mitochondria distributed along the cell; oligodendrocytes: more mitochondria concentrated on the cell soma and fewer, fragmented mitochondria in the proximal processes). Initially, we drew a circular ROI around the soma of the targeted cells. The area within the ROI was then irradiated with a 405 nm laser at 30% power and a speed of 200 Hz and scanned 60 times for a total of 77 seconds using the confocal microscope. Before and after images of the unconverted and photoconverted mito-Dendra2 were captured sequentially using the 488 nm and 552 nm laser wavelengths respectively. Repeated imaging was done over 4, 24, and 48 hours using consistent image acquisition settings. Image analysis was done using Fiji to follow changes in mitochondrial dynamics over time. The Z-stacks were smoothed, and a circular ROI was drawn around the soma of the targeted cell (replicating the ROI drawn for the photoconversion) on a single slice that showed strongest mito-Dendra2 signal. The same ROI was drawn for all the time points. The mean intensity within the ROI was measured for the unconverted (488 nm) and photoconverted (552 nm) mito-Dendra2 channels. The ratio of the latter to the former was taken for every time point and normalized to the ratio resulting from the time point after photoconversion.

### Mitochondria motility

To determine mitochondrial motility, we tracked the mitochondrial displacement over 30 minutes in awake, anesthetized (isoflurane or ketamine/xylazine), dexmedetomidine hydrochloride, or vehicle-treated animals. All images were acquired once a minute on the two-photon microscope from 2–3-month-old and 18-21-month-old animals. For awake imaging, the animals were allowed to adjust to the Mobile HomeCage for 30 minutes before imaging. Similarly, the imaging was done 30 minutes after isoflurane anesthesia (0.8-1%), 15 minutes after intraperitoneal injection of ketamine/xylazine (100 mgkg^-1^/10 mgkg^-1^), 30 minutes after intraperitoneal injection of 0.5 mgkg^-1^ dexmedetomidine hydrochloride (Tocris # 2749) dissolved in distilled water, and 30 minutes after intraperitoneal injection of vehicle. The animal’s velocity and locomotion pattern were tracked using the tracking software (version 3.0.0.65 / Beta1) provided with the Neurotar’s Mobile HomeCage.

To analyze mitochondrial motility, the time series were blinded, the cells of interest were cropped, and the Z-stacks were corrected for the 3-D drift with reference to the cytoplasmic channel. Z-projections spanning 15µm were created similarly for each condition, re-corrected for the 3-D drift with reference to the mitochondrial channel, smoothed, and auto-thresholded. The mitochondria coming from other cells as well as from the soma of the cell of interest were excluded from the analysis. Mitochondria were automatically tracked using the TrackMate^79^ (version 7.7.2) plugin in Fiji to determine the displacement of mitochondria tracks and instantaneous velocity. The images were filtered by applying a LoG detector with an estimated object diameter of 3 µm, quality threshold of 1 and the sub-pixel localization was checked. The Simple LAP Tracker was applied with a linking max distance of 3 µm, gap-closing max distance of 3 µm and max frame gap of 2. The tracks with less than 5 spots were excluded and the tracks with displacement of 3 µm or above were considered as motile. Post-hoc visual inspection was done to correct for major errors in particle tracking.

### Statistical analysis

All statistical analysis was performed using GraphPad Prism (version 9.5.1) or Excel. Each dot represents a cell, the lines/traces are at the mean and the error bars represent the s.e.m. Normal distribution was assumed for all the datasets and sample numbers were determined based on previous publications^62,66^. Animals were randomly assigned to each experimental group. The cells analyzed throughout the study were selected solely based on quality of the images and ability to distinguish individual cells and cell compartments. The cells analyzed for the mitochondria motility experiments were randomly selected with reference only to the first time point of the time series. The cells analyzed from the EM dataset were selected only based on what best satisfied the identification criteria for the cell types analyzed. Data was blinded for each condition of mitochondria motility analysis (awake, isoflurane, ketamine/xylazine, dexmedetomidine hydrochloride, vehicle) and for animal age when analyzing mitochondria length and distribution in young and aged animals. Throughout all the study, when defining ROIs of the cell compartments, the experimenter was blinded to the mitochondria channel and referred only to the cytoplasmic channel to prevent any bias on the regions selected. No animals were excluded from the statistical analysis.

**Supplementary Video 1: Mitochondria in an OPC that differentiates.** In vivo time series depicting a differentiating OPC and its mitochondria over 25 consecutive days. The mitochondria distribution begins to change with the start of myelin sheath formation at P79, initially increasing content in the periphery which then drops as the cell matures until it reaches a low-density steady state (3 frames per second).

**Supplementary Video 2: Mitochondria in an OPC that stays an OPC.** In vivo time series depicting an OPC and its mitochondria for 23 consecutive days where the fate of the OPC remains unchanged. Notice the overall consistent distribution of mitochondria in the cell over time (3 frames per second).

**Supplementary Video 3: Mitochondria in a dying OPC.** In vivo time series of 12 consecutive days depicting an OPC that undergoes cell death. The mitochondria content scales with changes in cell morphology before the formation of apoptotic bodies at P107 (3 frames per second).

**Supplementary Video 4: Mitochondria motility in an OPC.** Awake time series depicting dynamic mitochondria repositioning within an OPC over 30 minutes (10 frames per second, the video is set to loop three times).

**Supplementary Video 5: Stationary mitochondria in an oligodendrocyte.** Awake time series showing stationary mitochondria in an oligodendrocyte over 30 minutes (10 frames per second, the video is set to loop three times).

**Supplementary Video 6: Isoflurane anesthesia decreases OPC mitochondria motility.** Left: Still image of mitochondria in the cytoplasm of the target OPC at the start of the awake time series showing the initial distribution of mitochondria in the cell. Right: 30-minute time series showing higher mitochondria motility in awake than after isoflurane anesthesia (10 frames per second, the video is set to loop three times).

**Supplementary Video 7: Ketamine/xylazine anesthesia decreases OPC mitochondria motility.** Left: Still image of mitochondria in the cytoplasm of the target OPC at the start of the awake time series showing the initial distribution of mitochondria in the cell. Right: 30-minute time series showing higher mitochondria motility in awake than after ketamine/xylazine anesthesia (10 frames per second, the video is set to loop three times).

**Supplementary Video 8: Dexmedetomidine-induced sedation decreases OPC mitochondria motility.** Left: Still image of mitochondria in the cytoplasm of the target OPC at the start of the awake time series showing the initial distribution of mitochondria in the cell. Right: 30-minute time series showing higher mitochondria motility in awake than after administration of dexmedetomidine hydrochloride (10 frames per second, the video is set to loop three times).

**Supplementary Video 9: OPC mitochondria motility is comparable in control conditions.** Left: Still image of mitochondria in the cytoplasm of the target OPC at the start of the awake time series depicting the initial distribution of mitochondria in the cell. Right: 30-minute time series depicting similar motility rate of mitochondria in awake state and after vehicle-administration (10 frames per second, the video is set to loop three times).

**Supplementary Video 10: Reduced mitochondrial motility in aged OPCs.** Top: Still images of mitochondria in the cytoplasm of a target young (left) and aged (right) OPC at the start of the awake time series depicting the initial distribution of mitochondria in the cell. Bottom: 30-minute time series showing reduced mitochondria motility in aged OPC (10 frames per second, the video is set to loop three times).

